# Genetic mechanisms for impaired synaptic plasticity in schizophrenia revealed by computational modelling

**DOI:** 10.1101/2023.06.14.544920

**Authors:** Tuomo Mäaki-Marttunen, Kim T. Blackwell, Ibrahim Akkouh, Alexey Shadrin, Mathias Valstad, Tobjørn Elvsåashagen, Marja-Leena Linne, Srdjan Djurovic, Gaute T. Einevoll, Ole A. Andreassen

## Abstract

Schizophrenia phenotypes are suggestive of impaired cortical plasticity in the disease, but the mechanisms of these deficits are unknown. Genomic association studies have implicated a large number of genes that regulate neuromodulation and plasticity, indicating that the plasticity deficits have a genetic origin. Here, we used biochemically detailed computational modelling of post-synaptic plasticity to investigate how schizophrenia-associated genes regulate long-term potentiation (LTP) and depression (LTD). We combined our model with data from post-mortem mRNA expression studies (CommonMind gene-expression datasets) to assess the consequences of altered expression of plasticity-regulating genes for the amplitude of LTP and LTD. Our results show that the expression alterations observed *post mortem*, especially those in anterior cingulate cortex, lead to impaired PKA-pathway-mediated LTP in synapses containing GluR1 receptors. We validated these findings using a genotyped EEG dataset where polygenic risk scores for synaptic and ion channel-encoding genes as well as modulation of visual evoked potentials (VEP) were determined for 286 healthy controls. Our results provide a possible genetic mechanism for plasticity impairments in schizophrenia, which can lead to improved understanding and, ultimately, treatment of the disorder.

## 1 Introduction

Schizophrenia (SCZ) is a severe mental disorder, characterized by negative, positive and cognitive symptoms [Jauhar et al., 2022]. Although some medications have proved successful in constraining the positive symptoms, few drugs help improve the negative and cognitive symptoms of SCZ [McCutcheon et al., 2023]. The mechanisms of SCZ symptoms are far from understood but it is believed that they involve impaired function at the cellular level. Previous studies have suggested synaptic plasticity, which is a key cellular mechanism behind cognitive functions such as learning and memory, as one of the overarching cellular-level abnormalities in the disorder pathology [Stephan et al., 2006, Mould et al., 2021, Hall and Bray, 2022, Zhang et al., 2022].

Synaptic plasticity depends on the orchestrated action of a multitude of second messenger molecules and proteins. Genome-wide association studies (GWASs) have identified variants associated with risk of SCZ in hundreds of genes [Ripke et al., 2014, Trubetskoy et al., 2022], and many of these genes are fundamental for neuromodulation and synaptic plasticity [Devor et al., 2017]. Conclusive experimental data on the effects of SCZ-associated variants on plasticity mechanisms is difficult to obtain due to the large number of implicated gene variants. In fact, apart from a few ultra-high risk variants that have been studied in genetic animal models, it is not known how the variants affect plasticity either in a single neuron or at a network level. Here, we address this question by means of computational modelling of biochemical signalling pathways. Computational models provide an efficient means of assessing synaptic plasticity and exploring how it is affected by variations in the patients’ genetic make-up. We used the CommonMind gene-expression datasets from post-mortem brains of SCZ patients and healthy controls (HCs) [Hoffman et al., 2019] that provide the information on whether the expression of genes, particularly those associated with risk single-nucleotide polymorphisms (SNPs), are up-or down-regulated or unaffected in SCZ. We applied a computational model [Måaki-Marttunen et al., 2020] to the patients’ brain gene expression data and investigated the effects of gene variants on the pathways that lead to long-term potentiation (LTP) and depression (LTD). In this way we could unravel how genetic variants associated with SCZ relate to central synaptic plasticity mechanisms.

Although SCZ symptoms are difficult to study due to a lack of suitable corresponding conditions in animals *in vivo*, there are other robust phenotypes of SCZ that are quantifiable via electrophysiology [Wang et al., 2022]. A mental disorder phenotype strongly suggestive of a deficit in synaptic plasticity is the modulation of visual evoked potential (VEP), a.k.a. LTP-like plasticity of VEP, which is impaired in SCZ patients compared to HCs [Çavus et al., 2012]. A specific aim of the current study was to explore how this phenotype is altered by variants of SCZ-associated genes.

Using the model of [Måaki-Marttunen et al., 2020] we explored the impact that the altered expression profile of plasticity-regulating genes, as measured in prefrontal cortices (PFC) and anterior cingulate cortices (ACC) of SCZ patients, have on several forms of synaptic plasticity. We showed that both the conventional forms of plasticity and the plasticity induced by the VEP protocol are significantly impaired by the alterations of protein levels suggested by post-mortem expression data and that these alterations are driven by impaired PKA-pathway-mediated synaptic potentiation. We used a genotyped data set from a VEP modulation experiment of 286 HCs to validate our results and found that the associations between polygenic risk scores (PRSs) of SCZ based on groups of genes relevant for synaptic plasticity supported our predictions for the impairments of synaptic plasticity caused by altered gene expression. Our results uncover mechanisms behind the phenotype of altered VEP modulation and help to narrow down the pathways through which cortical synaptic plasticity is impaired in SCZ.

## 2 Results

### 2.1 The model reproduces numerous forms of plasticity

Synaptic plasticity in the cortex is dependent on a wide set of molecular mechanisms that vary between cortical areas and stages of development. To investigate the role of diverse pathways in different forms of plasticity affected by SCZ, we enhanced our model [Måaki-Marttunen et al., 2020] to include neurogranin, which is critical for plasticity [Zhong et al., 2009], and improved the model of GluR1 cycling (see Methods). We then validated our model (illustrated in Fig. 1A) by fitting the parameters to different data sets using multi-objective optimisation where the quantity of different proteins were varied to match the LTP/LTD outcome observed in the experiments. We were able to reproduce all the data as in the original model [Måaki-Marttunen et al., 2020] and found parameter sets that reproduced CaMKII-and stimulation frequency-dependent plasticity in V1 [Kirkwood et al., 1997] in adult (Fig. S1A) and young (Fig. S1B) animals. We also successfully reproduced the effects of PKC-neurogranin interactions on postsynaptic LTP in CA1 [Zhong et al., 2011] (Fig. S1C) and *β*-adrenergic receptor and stimulation frequency-dependent plasticity in V1 [Fischer et al., 2004] (Fig. S1D).

**Figure 1:**
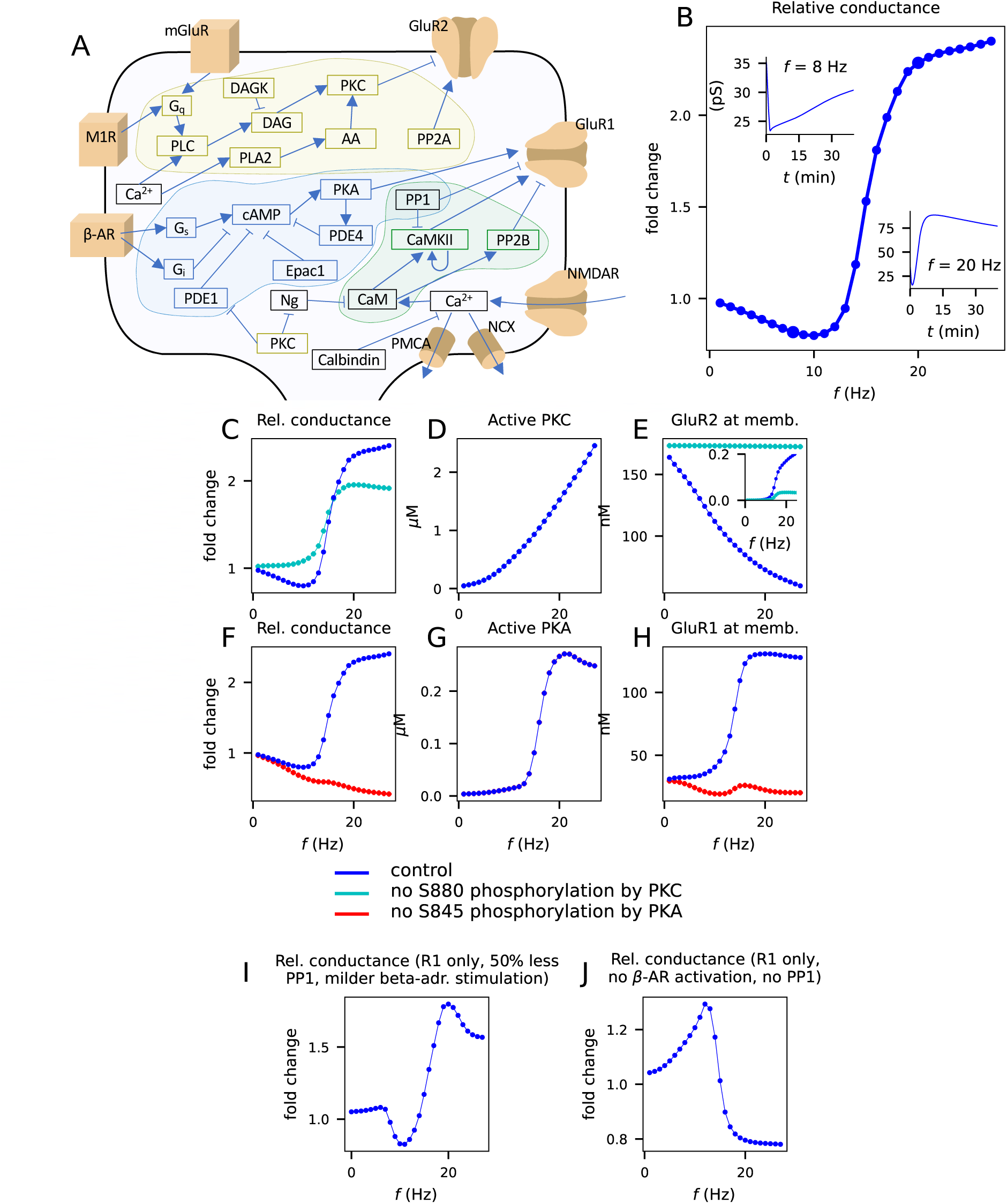
The model predicts Bienenstock-Cooper-Munro (BCM)-like plasticity where small input frequencies induce LTD and large ones LTP. **A:** Illustration of the cell membrane-level mechanisms and intracellular signalling pathways described by our model. **B:** Relative synaptic conductance 30 min after stimulus onset as a function of input frequency, given a fixed stimulation time of T=100 s. Insets: Time course of the synaptic conductance for input frequencies 8 Hz (left) and 20 Hz (right). **C:** Relative synaptic conductance 30 min after stimulus onset in the control case (blue) and when phosphorylation of S880 of GluR2 by PKC was disabled (cyan). **D:** Peak concentration of active PKC (all forms) during the simulation as a function of input frequency. **E:** Concentration of membrane-inserted GluR2 subunits 30 min after stimulus onset in the control (blue) and S880-blocked (cyan) cases. Inset: Probability of membrane-inserted AMPAR tetramers being homomeric R1-type 30 min after stimulus onset according to our statistical rule (Eq. 1; see [Måaki-Marttunen et al., 2020] for details). **F:** Relative synaptic conductance 30 min after stimulus onset in the control case (blue) and when phosphorylation of S845 of GluR1 by PKA was disabled (red). **G:** Peak concentration of catalytic subunit of PKA during the simulation as a function of input frequency. **H:** Concentration of membrane-inserted GluR1 subunits 30 min after stimulus onset in the control (blue) and S845-blocked (red) cases. **I:** Relative synaptic conductance in a PP1-reduced, GluR1-only synapse with milder *β*-adrenergic stimulation. This synapse produces bidirectional plasticity that is independent of the PKC-GluR2 pathway. **J:** Relative synaptic conductance in a GluR1-only synapse without PP1 and without *β*-adrenergic stimulation. This synapse produces an inverse LTP/LTD curve. In the simulations of blocked S845 phosphorylation in panels E and G, the loss of S845 activity was compensated by a spontaneous phosphorylation where the reaction rate of the added reaction was fitted so that the baseline synaptic conductance was equal to that in the default model.

To compare the effects of SCZ for a range of plasticity protocols across brain regions, we used our unified model of post-synaptic plasticity with a standardized frequency-controlled stimulation protocol with which to study the effects of SCZ-associated genes on different types of plasticity. The stimulation protocol consisted of evenly spaced 3-ms Ca^2+^ input pulses for 100 seconds where we varied the frequency (and number) of the input pulses. We then measured the total synaptic conductance 30 min after the stimulus onset as a function of the stimulation frequency.

Using the default protein concentrations and a GluR1-to-GluR2 ratio of 1:3, our model produces a Bienenstock-Cooper-Munro (BCM) type plasticity curve, where low stimulation frequencies produced LTD and high stimulation frequencies produced LTP (Fig. 1B). The LTD was PKC dependent (Fig. 1C), mediated by GluR2 endocytosis (Fig. 1D–E), while the LTP induced by high stimulation frequencies was PKA dependent (Fig. 1F) and was mediated by GluR1 exocytosis (Fig. 1G–H). Moreover, the LTP induced by high stimulation frequencies was amplified by PKC (Fig. 1C) because the PKC-mediated GluR2 endocytosis increases the likelihood of AMPAR tetramers being large-conductance homomeric AMPARs when the stimulus frequency is high (Fig. 1E, inset) [Måaki-Marttunen et al., 2020].

The bidirectional plasticity of Fig. 1B relies on both GluR1 and GluR2 dynamics. However, purely GluR1-based bidirectional plasticity where the LTD was mediated by dephosphorylation of S845 has been observed in CA1 [Lee et al., 2010] and modelled in [Castellani et al., 2005]. When we used a GluR1-only synapse (GluR1-to-GluR2 ratio of 1:0) and decreased the PP1 concentration by 50%, our model predicted a BCM-like plasticity curve (Fig. 1I). Our model also predicted that when PP1 was removed and the *β*-adrenergic receptor activation was blocked the GluR1-only synapse switched into an anti-BCM or anti-Hebbian mode of plasticity (Fig. 1J) occasionally observed in the cortex [Letzkus et al., 2006] or striatum [Fino et al., 2005].

Taken together, our model equipped with default protein concentrations produces a stereotypic bimodal, BCM-like plasticity. Next, we use this framework for assessing the effects of SCZ-associated genes on cortical synaptic plasticity.

### 2.2 Risk genes encoding synaptic intracellular signalling proteins affect cortical LTP/LTD

Impaired synaptic plasticity has been suggested as a unifying mechanism behind SCZ symptoms and pathogenesis [Stephan et al., 2006, Forsyth and Lewis, 2017], but it remains unknown whether and how the implicated risk variants [Devor et al., 2017] can affect the induction of LTP/LTD. To identify the set of plasticity-related genes conferring a risk of SCZ, we used two sources: psychiatric GWAS data from the Psychiatric Genomics Consortium (PGC) and expression data from the CommonMind Consortium (see Methods). We used our unified model of post-synaptic plasticity to identify which alterations of expression of the risk proteins modify which types of cortical post-synaptic plasticity. We implemented the effects of SCZ genetic alterations by altering the expression (both over-and under-expression) of the implicated proteins by *±*20% and evaluated the plasticity curves in response to the standardized protocols described above.

The altered expression of the risk proteins identified by GWAS or expression data produced diverse effects on LTP/LTD in the cortex (Fig. 2). In the PKA pathway, overexpression of PP1 by 20% decreased the basal synaptic strength and amplified the LTP (Fig. 2A, red), and similar effects were obtained by underexpression of PKA (Fig. 2B, gray) and overexpression of PDE4 (Fig. 2C, red). These effects were mediated by altered S845-phosphorylation-driven membrane expression of GluR1 subunits (Fig. S2A–C,G–I). In the PKC pathway, overexpression of PLA2 amplified both LTD and LTP (Fig. 2D), similar to overexpression of PKC (Fig. S2M). The effects of these parameter changes are attributed to their impact on S880-phosphorylation-mediated GluR2 endocytosis (Fig. S2D–E,J–K). Altered expression of CaMKII by *±* 20% did not significantly alter the LTP/LTD amplitude (Fig. S2N). By contrast, overexpression of neurogranin (Fig. 2E) or NCX (Fig. 2F), both of which lead to an increased amount of input Ca^2+^ required to activate CaM, caused a rightward shift in the transition from LTD to LTP. Overexpression of NCX also decreased baseline I1 activity and, consequently, increased PP1 availability, which led to increased baseline GluR1 exocytosis (Fig. S2F) and thus an increased baseline conductance (Fig. 2F inset). Alterations of the expression of Gq (Fig. S2O), mGluR (Fig. S2P), calbindin (Fig. S2Q), DAGK (Fig. S2R), or Gi (Fig. S2S) by *±* 20% had little or no effect. Decreased expression levels (gray) of the risk proteins always had qualitatively opposite effects than the increased levels (red; Fig. 2A–F, Fig. S2M–S).

**Figure 2:**
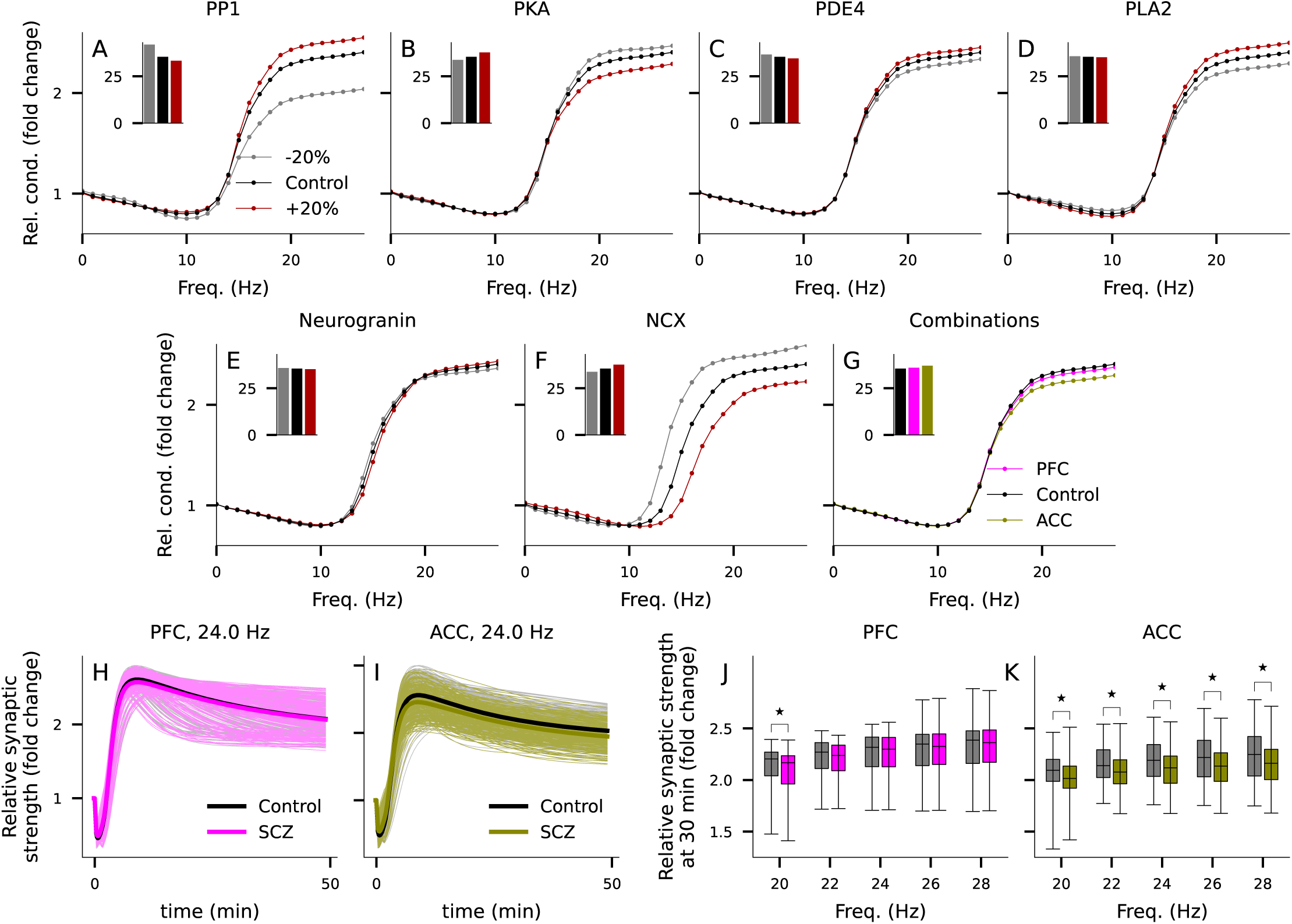
Altered expression of synaptic proteins associated with the risk of SCZ can impair or enhance LTP/LTD. **A–F**: The experiment of Fig. 1B was repeated with altered expression of the risk proteins of Table 1A, namely PP1 (A), PKA (B), PDE4 (C), PLA2 (D), neurogranin (E), and NCX (F). Red and gray curves represent the data from simulations where the underlying protein concentration was 20% higher (red) or lower (gray) than in the default model (black). The x and y axes show the frequency of inputs and the relative synaptic conductance 30 min after stimulus onset, respectively. Insets: the baseline synaptic conductance (pS). **G**: The experiment of Fig. 1B was repeated with a combination of expression-level alterations of the 10 genes affecting 9 model parameters (for PFC, pink) or the 12 genes affecting 9 model parameters (for ACC, green; see Table 1A) as suggested by post-mortem data. **H–I**: The experiments of Fig. 2G were repeated using subject-wise expression patterns for the underlying genes. The plots show the time courses of the relative synaptic conductance in response to 24 Hz stimulation for 100 s. The dim gray curves represent the subject-wise simulation results for HC subjects (N=215 in PFC and N=251 in ACC) and the dim pink (PFC; N=211) and light green (ACC; N=227) curves represent those for SCZ subjects. The thick curves represent the averages over the diagnostic groups. **J–K**: Box plots of the predicted post-30 min synapse weights for stimulation frequencies 20–28 Hz in the subject-wise simulations. The asterisks denote statistically significant differences between the diagnostic groups (U-test, p*<*0.05).

**Table 1:**
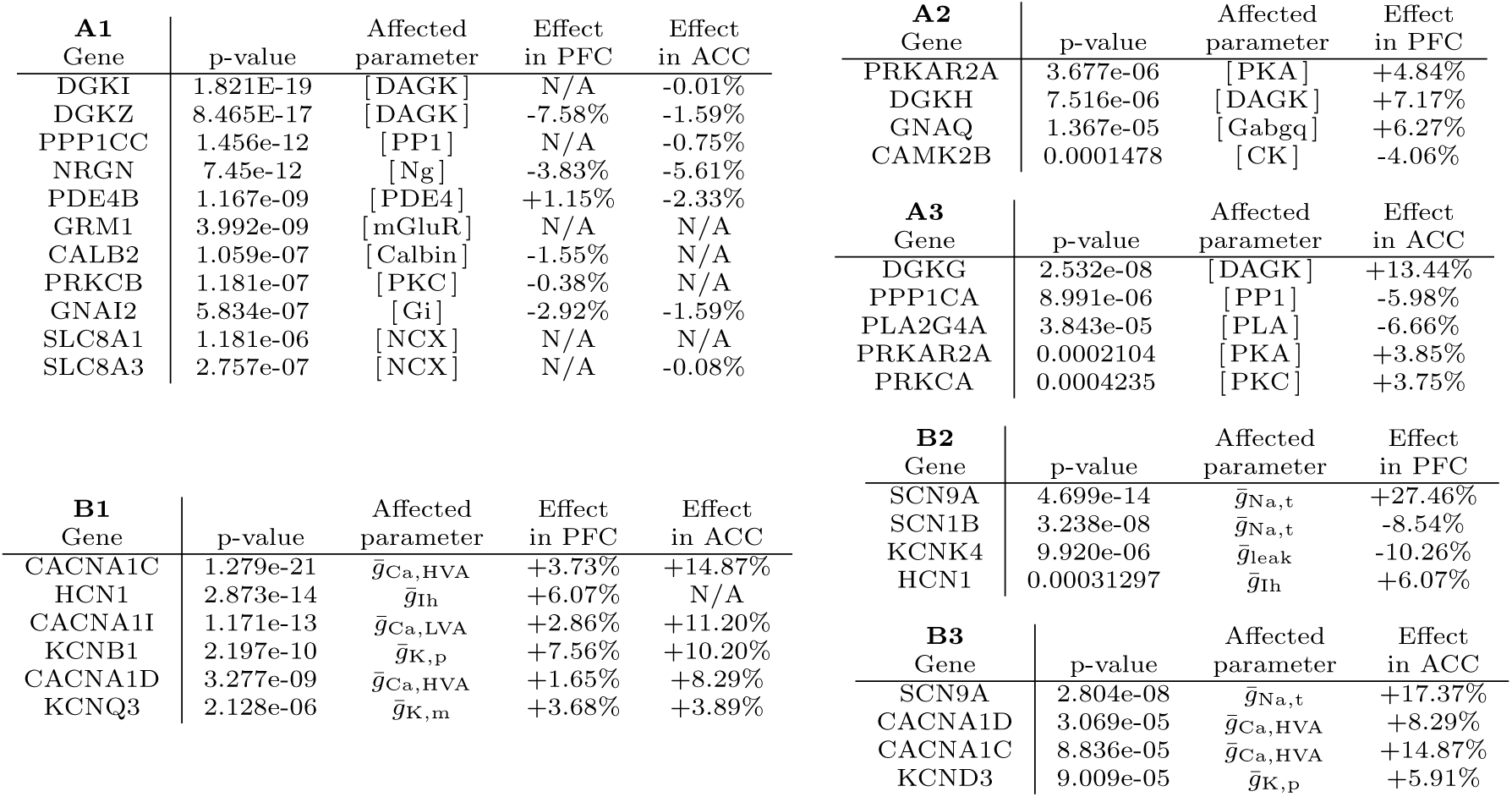
Table of synaptic (A) or ion channel-encoding (B) genes associated with SCZ.

While the alterations in the gene-expression levels are likely to vary within SCZ patients due to the heterogenic disease phenotype and polygenic architecture, indications of the direction and magnitude of the altered expression can be obtained from post-mortem studies. We estimated the relative difference in the expression levels of the SCZ-associated genes of Table 1A between SCZ patients and HCs in PFC and ACC using the CommonMind data. We repeated the simulations of Fig. 2A–F and Fig. S2M–S with these expression-level factors, either all combined (Fig. 2G) or a single protein-concentration change at a time (Fig. S3A–K). When all expression alterations were combined, both PFC-and ACC-like alterations affected the baseline synaptic strength and the LTP magnitude, but the effects of the variants corresponding to expression data from PFC were milder (LTP amplitude decreased by 2.2%) than the effects of those from ACC (LTP amplitude decreased by 8.0%; Fig. 2G). The predicted alterations of LTP magnitude in the PFC were mostly driven by altered PKA expression, whereas those in the ACC were mostly driven by altered expression of PKA, PP1, and PLA2 (Fig. S3A–C): these alterations caused the LTP amplitude to be decreased by 1.4% (PKA alteration in ACC, Fig. S3A), 3.3% (PP1 alteration in ACC, Fig. S3B), 2.5% (PLA2 alteration in ACC Fig. S3C), or 1.7% (PKA alteration in PFC, Fig. S3A) while alterations of other risk proteins affected the LTP amplitude by -0.5–0.8% (Fig. S3D–K).

To confirm our finding of impaired LTP, we simulated the plasticity outcome using a personalized approach, where the initial concentrations of proteins of Table 1A were determined in a subject-wise manner based on the CommonMind expression data (Fig. 2H–I). The amplitude of the plasticity was significantly (U-test, p*<*0.05) decreased in the SCZ group for the LTP induced by 20, 22, 24, 26, and 28 Hz stimulation in the simulations based on ACC expression data (p=2.5*·*10*^−^*^5^, 7.2*·*10*^−^*^6^, 9.1*·*10*^−^*^6^, 6.1*·*10*^−^*^5^, and 0.00016 for 20, 22, 24, 26, and 28 Hz, respectively), but only the decrease in LTP induced by 20 Hz stimulation was significant (p=0.0058 for 20 Hz, whereas p=0.067, 0.59, 0.73, and 0.80 for 22, 24, 26, and 28 Hz, respectively) in the simulations based on the PFC expression data (Fig. 2J–K). We also confirmed the robustness of this result by simulating the plasticity in synapses where the expression of the risk genes was altered using alternative strategies for determining the parameter changes based on the CommonMind data. A majority of these strategies yielded similar result, namely, that the gene-expression profiles of SCZ subjects compared to HCs in post-mortem ACC predicted an impaired LTP amplitude (Fig. S4).

Taken together, our simulations show that mild changes in expression levels of proteins associated with the risk of SCZ can weaken or strengthen the cortical LTP/LTD, and that the expression-level alterations observed *post mortem* in ACC of SCZ patients can cause a significant decrease in LTP amplitude.

### 2.3 Risk genes encoding both voltage-gated ion channels and intracellular signalling proteins regulate cortical STDP

STDP is crucially dependent on the membrane excitability properties of the post-synaptic neuron. We have previously studied the effects of SCZ-associated ion channel-encoding genes on pyramidal cell excitability [Måaki-Marttunen et al., 2016, Måaki-Marttunen et al., 2017, Måaki-Marttunen et al., 2019a] using biophysically detailed neuron models, and the same framework can be used to investigate how the integration of STDP-inducing pre-and postsynaptic stimuli is affected by these genes. Here, we test the effects of alterations of ion-channel expression in the post-synaptic neuron on STDP and how they compare to the modifications to the STDP curve caused by altered expression of intracellular signalling proteins.

For this multi-scale investigation, we used 1) the biophysically detailed model for layer II/III pyramidal cell [Markram et al., 2015] (L23PC) for determining the size and time course of NMDAR-mediated Ca^2+^ inputs for each pre/post-synaptic pairing interval (Fig. 3A) and 2) our biochemically detailed model of signalling pathways mediating plasticity in response to Ca^2+^ and neuromodulatory inputs (see Methods). We altered the maximal ion-channel conductances of the risk proteins in the pyramidal cell model as suggested by the genetic analysis (see Table 1B). We also evaluated the interaction between altered expression of ion channel-encoding and plasticity-regulating proteins.

**Figure 3:**
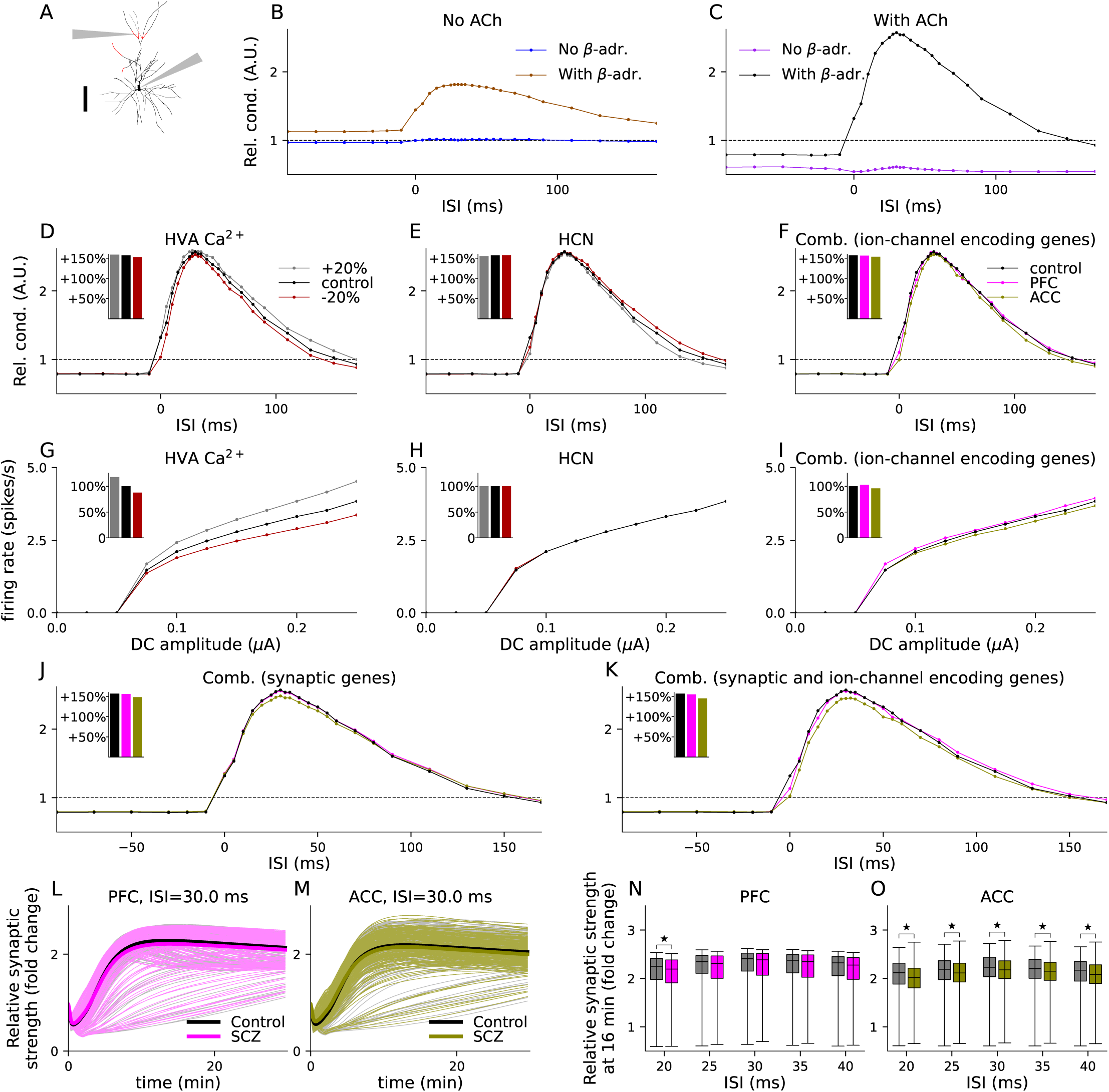
Altered expression of SCZ risk genes encoding voltage-gated ion channels or synaptic proteins of the post-synaptic neuron can affect STDP. **A:** Illustration of the L23PC model used for determining the Ca^2+^ time course. Compartments where the post-synaptic spine can be inserted are highlighted in red. The top arrow represents an exemplary location of the post-synaptic spine and the bottom arrow points at the soma where the burst of square pulse currents is injected. Scale bar 100 µm. **B–C:** Our model reproduces neuromodulator-gated STDP reported in [Seol et al., 2007]. The curves show the relative synaptic strength in response to 120 stimuli 16 minutes after stimulus onset as a function of inter-stimulus interval (negative values of ISI correspond to post-pre stimulus protocols) in the absence cholinergic and *β*-adrenergic neuromodulation (B, blue), in the absence cholinergic but presence of *β*-adrenergic neuromodulation (B, brown), in the absence *β*-adrenergic but presence of cholinergic neuromodulation (C, purple), and in the presence of both neuromodulators (C, black). **D–E:** The STDP curve when the post-synaptic neuron expresses 20% more (red) or less (gray) voltage-gated ion channels, namely, high-voltage-activated Ca^2+^ channels (D) or *I_h_* channels (E). Insets: maximal LTP value of the curve. **F:** The STDP curve with the PFC-(pink) and ACC-like (green) combinations of variants of ion-channel encoding genes. **G–I:** The f-I curves, i.e., spiking frequency of the post-synaptic neuron as a function of the amplitude of DC applied to soma, corresponding to the variants of panels D–F. Insets: The firing rate in response to DC of amplitude 0.25 µA, normalized by the response of the default neuron model (black). **J:** The STDP curve for the combinations of variants of synaptic risk genes as in Fig. 2G. K: The STDP curve for the combinations of synaptic protein-affecting variants of panel J and ion-channel-affecting variants of panel F. **L–M:** The time course of the relative synaptic strength in subject-wise simulations of LTP induction in response to pre-post stimuli with 30 ms interval. The dim gray curves represent the subject-wise simulation results for HC subjects and the dim pink (PFC; L) and green (ACC; M) curves represent those for SCZ subjects. The thick curves represent the averages over the diagnostic groups. N–O: Box plots of the predicted post-15 min synapse weights in PFC (N) and ACC (O) in the subject-wise simulations of pre-post intervals 20–40 ms. The asterisks denote statistically significant differences between the diagnostic groups (U-test, p*<*0.05). simulating VEP-plasticity protocol and tested the effects of SCZ-associated genes on this form of plasticity.

First, we validated our updated model by replicating the experimentally observed neuromodulator-gated STDP curve for layer IV*→*II/III pyramidal cell synapses in the visual cortex. Similar to experimental data [Seol et al., 2007] and our previous model [Måaki-Marttunen et al., 2020], little or no plasticity is observed in the absence of neuromodulators (Fig. 3B, blue), LTP is obtained when the pairing stimuli are administered with *β*-adrenergic input (Fig. 3B, brown), LTD is obtained when the stimuli are administered with cholinergic input (Fig. 3C, purple), and pairing-interval dependent LTP/LTD is obtained when both neuromodulators are present (Fig. 3C, black). Next, we simulated the effects of altered expression of ion channels in the post-synaptic neuron on the cortical STDP (Fig. 3D–E; Fig. S5A–E). Overexpression of CACNA1C (Fig. 3D, red) and KCNQ3 (Fig. S5A) led to decreased LTP amplitudes — a mild decrease of LTP amplitude was also observed for increased leak channel conductance (a putative effect of overexpression of KCNK4; Fig. S5B). By contrast, overexpression of HCN1 mildly strengthened LTP (Fig. 3E). The increase of LTP amplitude by increase of HCN channels is explained by a generally increased dendritic excitability [Måaki-Marttunen and Måaki-Marttunen, 2022], while the LTP-impairing effect of increased high-voltage activated (HVA) Ca^2+^ conductance was caused by increased SK activity [Måaki-Marttunen et al., 2017]. Altered expression of SCN1B or SCN9A (Fig. S5C), KCNB1 or KCND3 (Fig. S5D), and CACNA1I (Fig. S5E) had little effect on STDP. The effects of the altered expression of ion channels on post-pre LTD were small (Fig. 3D–E; Fig. S5A–E). We also analysed the effects of these variants on neuron excitability by measuring the predicted f-I curves, namely, firing frequencies in response to DC of different amplitudes. The maximal LTP amplitudes of the *±*20% variants were correlated (Pearson correlation coefficient 0.71, p-value = 0.0041) with the firing frequencies in response to somatic DC, indicating that the increase in LTP was driven by a generic increase in the single-cell excitability (Fig. 3G–H; Fig. S5F–J). Exceptions to this pattern were the SCN1B or SCN9A-encoded fast Na^+^ channel, where overexpression had little effect on LTP response but increased the firing response (Fig. S5C,H), and HCN and M-type channels where altered expression affected the STDP but had little or no effect on somatically induced firing of action potentials (Fig. 3E,H; Fig. S5A,F).

Simulations with combinations of altered expression levels of these voltage-gated ion channels as measured in PFC and ACC showed that LTP amplitude may be reduced (Fig. 3F) and intrinsic pyramidal cell excitability altered (Fig. 3I) in PFC and ACC in SCZ patients. The effects on LTP amplitude were, however, mild (maximal LTP amplitude +156% and +154% in PFC-and ACC-like alterations, respectively, compared to +157% in control; Fig. 3F) suggesting that the changes in STDP caused by expression-level alterations of different ion channels can compensate for each other. The effects on intrinsic pyramidal cell excitability were different for PFC-like alterations compared to ACC-like alterations (firing frequency increased by 2.7% and decreased by 4.1% for PFC-and ACC-like alterations, respectively; Fig. 3I).

We asked how the alterations in ion channel-encoding and plasticity regulating genes interact to affect the STDP. We first simulated the effects of altered expression of intracellular signalling proteins (Fig. 3J). The effect of PFC-like alterations was smaller (maximal LTP amplitude +156% compared to +157% in control) than those of ACC-like alterations (maximal LTP amplitude +148%; Fig. 3J inset). When the combination of the ion-channel variants was used together with the combination of synaptic gene variants, the LTP amplitude was more strongly decreased, particularly in a synapse expressing ACC-like alterations (maximal LTP amplitude +155% and +145% in PFC-and ACC-like alterations, respectively, compared to +157% in control; Fig. 3K). We confirmed the significance of the effects of the ACC-like alterations by performing subject-wise simulations (Fig. 3L–M). The amplitude of the LTP induced by pre-post pairs of stimulus (ISI=20, 25, 30, 35, 40 ms) stimuli was significantly smaller (U-test, p*<*0.05) in the SCZ subjects than in the HCs according to the ACC post-mortem expression data (p=0.0026, 0.018, 0.045, 0.038, 0.021 for ISI=20, 25, 30, 35, and 40 ms, respectively; Fig. 3O), but the corresponding difference in simulations based on PFC expression data was only significant for 20 ms ISI (p=0.035, 0.093, 0.23, 0.16, and 0.11 for ISI=20, 25, 30, 35, and 40 ms; Fig. 3N).

Taken together, our results suggest that altered expression of ion-channel encoding SCZ-associated genes can alter the LTP amplitude in the STDP protocol, and simulations using post-mortem expression data, especially those from ACC, support a decrease in pre-post stimulus-induced LTP amplitude.

### 2.4 Plasticity in response to stimuli used in the clinical phenotype of VEP modulation may be altered by SCZ-associated gene variants

SCZ has been linked to synaptic abnormalities both by genetics analyses and post-mortem studies, but mechanistic links mapping these deviances to disorder symptoms and phenotypes is missing [Obi-Nagata et al., 2019]. A promising phenotype for study is visually evoked potentials, the plasticity of which is reported to be impaired in psychotic disorder patients [Normann et al., 2007, Ç avus et al., 2012]. The VEP is a visual stimulus-related EEG response and plasticity or modulation of VEP is the potentiation of VEP amplitudes after long-lasting checkerboard reversal stimulation [Teyler et al., 2005]. Although the LTP-like plasticity of VEP is likely to reflect synaptic plasticity occurring in multiple parts of VEP-related brain circuitry, it shares properties with traditional monosynaptic LTP, such as NMDAR-dependency [Normann et al., 2007, Cooke and Bear, 2012, Clapp et al., 2012]. Here, we explored the cortical synaptic plasticity caused by stimuli

To model the synaptic inputs in the VEP protocol, we stimulated the synapse with an action potential every 500 ms to mimic a checkerboard reversal with a frequency of 2 reversals/s for 10 min [Zak et al., 2018], and measured the effects on synaptic conductance 10–60 min after the stimulus onset. Each synaptic input was modelled as a square pulse influx of Ca^2+^ (150 particles/ms) lasting for 3 ms, which was by default accompanied by a flux of *β*-adrenergic agonist (5 particles/ms for 3 ms) as well as glutamate and cholinergic ligands (10 particles/ms for 3 ms) into the vicinity of the synapse, as in the experiments of Fig. 1–2. We tested the LTP/LTD response of the synapse to these stimuli using two extreme GluR1:GluR2 ratios, namely, 1:0 (GluR1-type) and 0:1 (GluR2-type).

Our model predicted that both GluR1-type (LTP; Fig. 4A–B) and GluR2-type (LTD; Fig. 4C–D) synapses underwent plasticity that was maintained for more than 1 hour. Neuromodulators were critical, as in the absence of neuromodulators the VEP protocol did not induce long-lasting plasticity changes (Fig. 4A,C).

**Figure 4:**
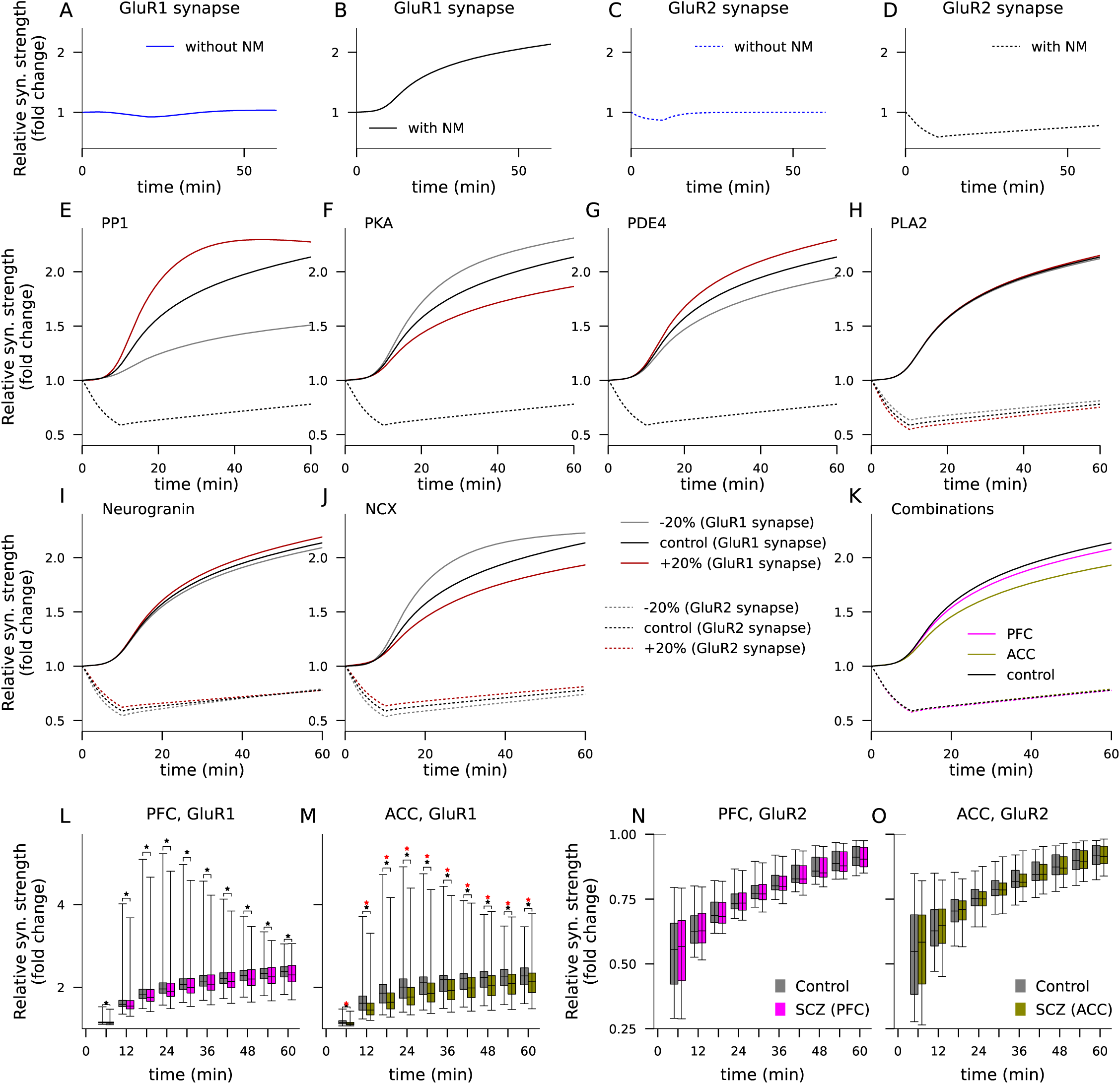
Altered expression of risk genes encoding synaptic proteins alters the plasticity response to VEP-like stimulus protocols. **A–D**: Time course of relative synaptic conductance in response to VEP-like stimulus (1200 stimuli administered during the 0–10 min interval) in the absence (A,C; blue) or presence (B,D; black) of neuromodulators in a synapse where all GluRs were GluR1 (A–B; solid) or GluR2 (C–D; dashed) type. **E–J**: Time course of the synaptic conductance in the VEP protocol with neuromodulatory stimulation in GluR1 (solid) or GluR2-type (dashed) synapses when the expression of SCZ risk protein PP1 (E), PKA (F), PDE4 (G), PLA2 (H), neurogranin (I), or NCX (J) was altered. **K**: Time course of the synaptic conductance for a combination of PFC-like (pink) and ACC-like (green) variants. **L–O**: The experiments of panel K were repeated using subject-wise expression patterns. The box plots show the relative synaptic conductance in response to VEP-like stimulus in a GluR1-type (L–M) or GluR2-type (N–O) synapse. Data for 6–60 min after stimulus onset are shown for both control (gray) and SCZ (pink, green) population for PFC-(L,N) and ACC-based (M,O) simulations. The black asterisks denote statistically significant differences between the diagnostic groups (U-test, p*<*0.05), and the red asterisks denote differences that were significant using a stricted threshold (p*<*0.005).

Similar to the plasticity induced by the standard protocol, VEP plasticity was affected by alterations in PP1 (Fig. 4E), PKA (Fig. 4F), and PDE4 (Fig. 4G) in GluR1-type synapses, though not in GluR2-type synapses. In contrast, alterations of PLA2 (Fig. 4H) and PKC (Fig. S6A) affected the plasticity in a GluR2-type synapse but not GluR1-type synapse. Neurogranin and NCX affected the plasticity of both types of synapses (Fig. 4I–J). No other risk proteins (including CaMKII) modified plasticity in response to the VEP-like stimulus when quantity was changed by *±*20% (Fig. S6B–G). PFC-like alterations of expression levels only mildly reduced the plasticity in a GluR1-type synapse (the increase in synaptic strength at 60 min was +108% compared to +114% in control) but not the GluR2-type synapse (Fig. 4K, pink). By contrast, ACC-like alterations strongly reduced the VEP-stimulus-induced plasticity in the GluR1-type synapse (in a GluR1-type synapse, the increase in synaptic strength was +93% compared to +114% in control) but not the GluR2-type synapse (Fig. 4K, green). The effects of the variant combinations on the plasticity in the GluR1-type synapse were driven by modifications of PKA-pathway genes: the impacts of the variant combinations were nearly abolished in a GluR1-type synapse when the PKA-pathway-affecting genes were excluded from the combination (Fig. S6H–M).

We confirmed the significance of these findings with the simulations based on subject-wise expression data. In subject-wise simulations of the VEP-like stimulation of a GluR1 synapse, both PFC-and ACC-like expression alterations caused a significant decrease in the LTP amplitude for each considered time point (6–60 min after stimulus onset in 6-min intervals; U-test, p*<*0.05), but only the effects of the ACC-like expression alterations were significant when a smaller p-value threshold (U-test, p*<*0.005) was used (Fig. 4L–M). Consistent with the experiments based on group-averaged expression data, the differences between SCZ and HC in plasticity responses in the GluR2-type synapse were non-significant (Fig. 4N–O).

Taken together, our results suggest that the modulation of synaptic strength by a VEP-like stimulus protocol can be moderately altered by risk proteins of SCZ. When the expression of the risk proteins is altered to reflect the expression alterations observed *post mortem* in SCZ, the PKA-pathway-dependent potentiation of GluR1-containing synapses by VEP-like stimulus protocol is weakened. Our results shed light into the polygenic mechanisms of VEP modulation deficits in SCZ and may pinpoint a generic vulnerability of plasticity in the mental disorder.

### 2.5 Genotypes and electrophysiological data support the contribution of SCZ-associated plasticity-regulating genes to the deficit of VEP plasticity

Although our model predictions for effects of altered expression of SCZ on generic LTP/LTD are difficult to validate, our predictions for the effects on plasticity induced by VEP protocol can be compared to EEG data collected from corresponding experiments. Here, we tested whether our model predictions align with genetic associations found in the genotype-phenotype data. For this, we used a large data set from The Thematically Organised Psychosis (TOP) Study [Engh et al., 2010] containing genetic and EEG data of VEP modulation from 286 HCs [Valstad et al., 2020, Valstad et al., 2021]. We calculated PRSs based on SNPs in different plasticity-regulating or ion channel-encoding genes (see Methods and Table S4) for each subject and determined their association with the EEG phenotypes by fitting a linear model, where the EEG index was regressed against the considered PRS, age, and sex. We only included SNPs that were indicative of SCZ risk (p-value *<* 10*^−^*^5^) in the recent GWAS [Trubetskoy et al., 2022]. We quantified the associations using the Pearson correlation coefficient, the *β* coefficient and the p-value for the PRS obtained from the linear model.

Of the VEP indices measured 2–4 min after the end of the conditioning stimulation (12–14 min after the onset of the conditioning stimulation, 14–16 min after the first baseline stimulation), the modulation of C1 (Fig. 5A) and N1 (Fig. 5C) amplitude were not significantly associated with the PRSs that we considered, whereas the P1 modulation (Fig. 5B) was significantly associated with the following PRSs: 1) the PRS based on SNPs in the genes encoding ion channels and plasticity-related proteins of Table S4 (*β*=-141, correlation coefficient -0.12, p=0.039), 2) the PRS based on SNPs in a subset of plasticity-regulating genes affecting plasticity via PKA pathway (purple genes of Table S4; *β*=-68, correlation coefficient -0.13, p=0.029), and 3) the PRS based on SNPs in ion-channel encoding genes (light green genes of Table S4; *β*=-89, correlation coefficient -0.14, p=0.020). Fig. 5D,F,H plots the values of P1 modulation plotted against the tested PRS values. All these *β* coefficients were negative, indicating that the presence of SCZ-risk SNPs in these sets of genes were predictive of lower post-modulation P1 amplitude in the VEP experiment and thus suggestive of impaired LTP-like plasticity. These results held also for PRSs calculated from the same sets of genes using alternative thresholds (5*·*10*^−^*^6^ and 10*^−^*^6^) for the inclusion of the SNPs (Fig. S7A–B). We went on to confirm the relevance of the SNPs in synapse-specific genes by using an alternative set of genes, as introduced in [Lips et al., 2012]. Namely, we constructed a PRS based on the SNPs within the genes in the following categories (Table S4 of [Lips et al., 2012]): Intracellular Signal Transduction, Excitability, GPCR signalling, Protein cluster, Neurotransmitter metabolism, LGIC signalling, Ion balance/transport, and G-protein relay, totalling 432 genes. The modulation of P1 was significantly associated with this alternative synapse-specific PRS (*β*=-102, p=0.016; Fig. S7C).

**Figure 5:**
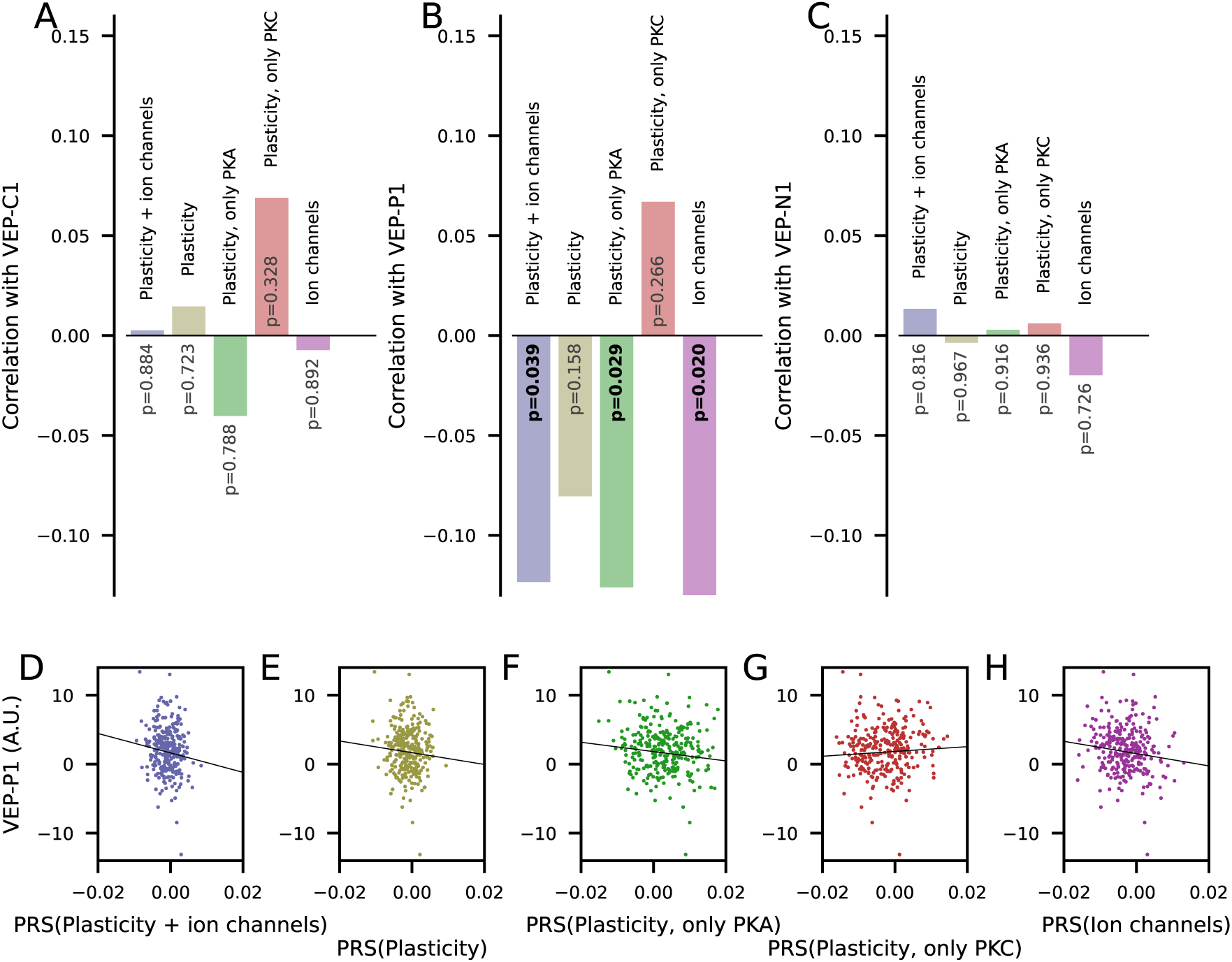
Analysis of a genotyped EEG dataset of VEP modulation provides support for the involvement of SCZ-associated variants of genes regulating synaptic plasticity and genes encoding ion-channel subunits in impairment of cortical plasticity. **A–C:** Pearson correlation coefficients between C1 amplitude (A), P1 amplitude (B), or N1b amplitude (C) and the five PRSs. **D–H:** P1 amplitude plotted against the five PRSs based on SNPs in the following gene sets: all genes of Table S4 (D), genes encoding plasticity-related proteins (E), genes encoding plasticity-related proteins that primarily couple to PKA (F) or PKC (G) pathway, or genes encoding ion channels or their subunits (H). The lines represent the effect of the PRS on the EEG index as obtained from the linear regression.

Taken together, our analysis suggests that the modulation of VEP-P1 is influenced by variants within plasticity-regulating and ion channel-encoding risk genes of SCZ. The directions of the associations suggest that the presence of SCZ risk SNPs in ion-channels-encoding or PKA-coupled genes regulating plasticity decreases the LTP-like plasticity of VEP-P1.

## 3 Discussion

In this work, we analysed the effects of SCZ-associated genes involved in synaptic plasticity on the amplitude and sensitivity of cortical LTP and LTD. We showed that mild alterations of expression levels of the proteins encoded by these genes can lead to significant alterations of threshold between LTD and LTP as well as LTP amplitude (Fig. 2), STDP amplitude (Fig. 3), and the plasticity induced by VEP-like protocols (Fig. 4). When we implemented the alterations of synaptic gene expression as observed in comparative post-mortem experiments, where gene expression in PFC and ACC were measured from SCZ patients and HCs, our model predicted a mild decrease in LTP amplitude for the PFC-like variant and a stronger decrease for the ACC-like variant (Fig. 2G). Similarly, both the PFC-like and ACC-like variants were predicted to reduce the amplitude of VEP-like plasticity, the ACC-like variant more than the PFC-like variant (Fig. 4N). The association of the genotype with the EEG response to VEP-like plasticity protocol (Fig. 5) supports our findings of impaired neocortical plasticity in SCZ.

There at least five different kinds of indirect experimental evidence of altered synaptic plasticity in SCZ (cf. [Forsyth and Lewis, 2017]). Post-mortem studies have reported 1) altered expression of synapse-associated genes [Fromer et al., 2016] and 2) aberrant spine morphology [Roberts et al., 1996] or spine density [Glausier and Lewis, 2013] in SCZ patients compared to HCs, the former observation being a likely culprit for altered synaptic plasticity in SCZ and the latter being its consequence. 3) Recent GWASes of SCZ have implicated a disproportionally large set of the genes that are associated with synaptic plasticity [Devor et al., 2017, Trubetskoy et al., 2022]. Furthermore, 4) certain phenotypes and endophenotypes of SCZ, such as altered modulation of LTP-like plasticity of VEP [Ç avus et al., 2012], impaired mismatch negativity [Garrido et al., 2009], and altered habituation of acoustic startle [Braff et al., 1978], together with 5) cognitive symptoms of SCZ such as impaired working memory [Lett et al., 2014] suggest that the very mechanisms of synaptic plasticity may be affected in the mental disease. Three of these modalities converge in this work. We selected the parameters to alter based on GWAS results (3), determined the magnitude of the parameter changes based on post-mortem expression studies (1), and predicted the effects on phenotypic measures (4) using stimulation protocols designed to mimic the synaptic inputs expected to occur in the experiments of these phenotypes. Furthermore, since many genetic manipulations that impair plasticity also impair working memory in animals [Borralleras et al., 2016, Chan et al., 2007], our results showing a decreased LTP amplitude may also prove relevant for deciphering the cause of cognitive dysfunction (5) in SCZ.

It should be noted that there is incomplete evidence on altered expression of GWAS-identified risk genes in SCZ, an assumption that our approach is partly based on. For example, Table 1 shows that while half of the considered ion channel-encoding GWAS-identified genes (CACNA1C, CACNA1D, and HCN1) reached the significance threshold in the expression data, none of the plasticity-regulating GWAS-identified genes did. Alternatives for this assumption are that the risk SNPs mediate the risk 1) by altering the transcription of other genes or 2) by direct effects on the structure of the encoded proteins. While usually only ultra-rare SCZ-associated variants are coding variants (i.e., result in proteins with mutated structure) [Harrison, 2015], the hypothesis that many SCZ-associated SNPs only regulate gene transcription of genes other than themselves cannot be discarded either for common or rare variants. Indeed, common SCZ-associated SNPs have been shown to alter gene methylation [Hannon et al., 2016, Jaffe et al., 2016], and some variants are located in expression quantitative trait loci (eQTLs) [Bhalala et al., 2018]. Nevertheless, previous data show that many GWAS-identified SCZ risk genes that have risk SNPs within their intra-gene region, e.g. CACNA1C, PDE4B, and NRGN (see Table 1), are themselves differentially expressed in SCZ [Yoshimizu et al., 2015, Fatemi et al., 2008, Broadbelt et al., 2006].

In this work, we specifically focused on a phenotype often encountered in SCZ patients and strongly suggestive of impairment of plasticity, namely, the deficit in the modulation of VEP [Çavus et al., 2012]. The animal-model correlate of this phenotype, the stimulus-specific response potentiation (SRP), has been found to be NMDAR dependent [Frenkel et al., 2006], making it likely to rely at least partly on post-synaptic plasticity. The SRP has been found obstructed by infusion of Myr-SIYRRGARRWRKL-OH (Myr-Zip) [Cooke and Bear, 2010], a peptide that strongly inhibits the atypical PKC isoform PKM-zeta but also incompletely inhibits PKC-alpha [Bogard and Tavalin, 2015]. Our model describes the PKC pathway alongside PKA and CaMKII pathways that are also central for cortical LTP/LTD and is thus well suited to analyze the effects of altered expression of many plasticity-associated proteins on VEP modulation.

It is not known what the circuit mechanisms behind VEPs and their modulation are, but one can safely hypothesize that the modulation-block-induced increase in the VEP amplitude is either due to potentiation of synapses positively contributing to or boosting the VEP amplitude or due to depression of synapses that restrict the VEP amplitude, or both. According to our model and most plasticity data from the cortex, potentiation is typically induced in synapses with GluR1-containing AMPARs and mediated by the PKA pathway, while depression is often induced in synapses with GluR2-containing AMPARs and mediated by the PKC pathway. Our modelling and data analysis showed that SCZ risk genes in the PKA pathway contribute to a statistically significant impairment of the VEP modulation (Fig. 4L–M, Fig. 5F) while the contribution of SCZ risk genes in the PKC pathway was ambiguous and non-significant (Fig. 4N–O, Fig. 5G). Our results thus suggest that the SCZ-associated deficit in VEP modulation is due to impairments in the former (potentiation of synapses that positively contribute to VEP-P1) rather than latter (depression of P1-hindering synapses) type of plasticity. A possible candidate for conveying this effect are L23PCs in visual cortex: they have been shown to undergo PKA-dependent LTP [Seol et al., 2007] and are a key population for controlling the cortical output [Quiquempoix et al., 2018] — they also detect a mismatch between visual and top-down inputs [Jordan and Keller, 2020], which may be relevant in stimulation protocols with abrupt changes such as the checkerboard reversal used in VEP. Substantial contribution to the modulation of VEP may also come from plasticity occurring in other cortical areas, including the ACC [Sidorov et al., 2020], one of the two source areas of our differential expression data in SCZ. Although we here tested only the effects of synaptic gene variants on the plasticity induced by the VEP-like protocols, our STDP modelling experiments with altered expression of ion channel-encoding genes (Fig. 3) and the association between the PRS based on ion channel-encoding genes and VEP modulation (Fig. 5H) suggest that SCZ-associated alterations in ion channel-encoding gene expression may impair the VEP modulation approximately as strongly as altered expression of synaptic plasticity-regulating genes.

Our results highlighting especially the PKA pathway in the predicted SCZ-associated alterations of plasticity help in narrowing down the generic hypothesis of impaired plasticity in SCZ. Recently, there has been increasing interest toward PDE4 inhibitors as a potential treatment of SCZ. Our results suggest that decreasing PDE4 activity increases the GluR1 membrane insertion and synaptic conductance at baseline and under mild but not strong Ca^2+^ inputs (Fig. S2C), leading to an occlusion of HFS-induced LTP in synapses expressing both GluR1 and GluR2 (Fig. 2K). Supporting data were obtained in [Wiescholleck and Manahan-Vaughan, 2012], where rolipram, a PDE4 inhibitor, increased the baseline synaptic strength without affecting the maximal synaptic strength attained in an HFS plasticity protocol (although it facilitated LTP in a longer time scale, *>* 2 hours after stimulus onset). Thus, the therapeutic effects of PDE4 inhibition may be mediated by increased baseline AMPAR activity and consequently increased basal excitability rather than improved plasticity mechanisms. This is supported by the finding that PDE4 expression is already reduced in SCZ patients [Fatemi et al., 2008]. Of note, other studies suggest that basal synaptic activity is normal or only mildly strengthened and that the amplitude of early LTP is also enhanced under PDE4 inhibition [Barad et al., 1998, Rutten et al., 2008]. However, even if PDE4 inhibition may be effective in restoring cognitive functioning in animals and patients where these phenomena are impaired, the small differences in the expression of PDE4 between SCZ and HCs suggest that the cause of the LTP deficits lies somewhere else than in an overexpression of PDE4.

Our model predictions were based on post-mortem RNA expression data from two brain regions (PFC and ACC data from the CommonMind), but our framework can be used with other expression data as well when they become available. In our PFC-and ACC-like variants, we included alterations of the concentration of many different proteins but the largest effects on the LTP amplitude were mediated by the expression levels of calcium-dependent intracellular PLA2 (encoded by PLA2G4A, under-expressed in post-mortem SCZ brains, see Table 1), RII-type PKA (encoded by PRKAR2A, over-expressed), and the catalytic subunit of PP1 (encoded by PPP1CA or PPP1CC, both under-expressed). Apart from PPP1CC, all these genes were identified using CommonMind data and had relatively low significance in the GWAS data (p-value *>* 10*^−^*^4^). Another plasticity-regulating gene that has previously been shown to be significant both in GWAS [Ripke et al., 2014, Devor et al., 2017] and expression [Broadbelt et al., 2006] is neurogranin-encoding NRGN. Contribution of variants of this gene to cognitive symptoms are supported by genetic studies that showed an association between SNPs of NRGN and cognitive abilities [Donohoe et al., 2010, Krug et al., 2013, Ohi et al., 2013]. Here, we showed that -20% change in neurogranin expression shifts the LTD/LTP curve leftwards (facilitated LTP) and +20% rightwards (hindered LTP; Fig. 2I). Overexpression of neurogranin was also accompanied by an increase in LTP amplitude for largest Ca^2+^ inputs (*>* 20 Hz stimulation in Fig. 2I) — this was in line with experimental data from CA1 [Hwang et al., 2021]. The CommonMind data sets predicted an expression difference of up to 6% only, which resulted in very small effects on the threshold and amplitude of LTP (Fig. S3G). However, much larger contribution of neurogranin to LTD/LTP could be expected since a decrease of 36–72% in neurogranin protein expression, measured by immunostaining, has been observed in post-mortem PFCs of SCZ patients [Broadbelt et al., 2006].

There currently exist no polygenic experimental models to have produced data that we could compare our results with, but our findings on effects of altered expression of single proteins can be confirmed with data from genetic and pharmacological experiments. A knock-out of NCX2 in mice caused a left-ward shift in the LTD/LTP curve in CA1 [Jeon et al., 2003] qualitatively similar to our 20% decrease in NCX concentration (Fig. 2J). Knock-out of a PP1 isoform, PP1*β*, increased basal synaptic transmission and attenuated LTP in CA1 [Foley et al., 2023] (N.B. currently unpublished) similar to our experiment of under-expression of PP1 (Fig. 2M), but opposite results were found for knock-out of other isoforms [Foley et al., 2023] and for pharmacological manipulation of PP1 [Wu et al., 2008, Morishita et al., 2001]. Pharmacological inhibition of PLA2 caused a decrease in LTP amplitude in CA1 [Massicotte et al., 1990], similar to our experiment of underexpression of PLA2 (Fig. 2G). As for proteins more thoroughly studied in plasticity experiments, particularly CaMKII and PKA, we have already shown that our model (both the previously published [Måaki-Marttunen et al., 2020] and the updated (Fig. S1) model) reproduces brain-area-specific data in the presence and absence of blockers or activators of these proteins.

The LTP/LTD pathways in our model are restricted to post-synaptic mechanisms, excluding structural plasticity as well as gene transcription and molecular trafficking between spine and soma. Among the CommonMind-identified genes were CAMK2G (p-value 3 *·* 10*^−^*^8^ in PFC and 1 *·* 10*^−^*^4^ in ACC), which encodes a CaMKII isoform that, once activated, delivers Ca^2+^-bound CaM into the nucleus [Ma et al., 2014] and has been hypothesized to regulate gene transcription [Ma et al., 2015], and CDK5 (p-value 1 *·* 10*^−^*^6^ in PFC and 3 *·* 10*^−^*^8^ in ACC), which encodes a cyclin-dependent kinase involved in up-regulation of GABAA-receptor activity and suppression of NMDA-receptor clustering [Cheung et al., 2006]. Furthermore, among the GWAS-identified genes [Trubetskoy et al., 2022] there are many that directly interact with proteins described in our model, such as the genes encoding protein-phosphatase-regulating subunits PPP1R13B, PPP1R16B, PPP2R2A, PPP2R2B, PPP2R3A, PPP2R3C, PPP2R4, PPP2R5B, and PPP3R1 (minimal p-value for SNPs in these genes were 6*·*10*^−^*^14^, 3*·*10*^−^*^16^, 8*·*10*^−^*^12^, 3*·*10*^−^*^6^, 2*·*10*^−^*^12^, 6*·*10*^−^*^7^, 1*·*10*^−^*^6^, 5*·*10*^−^*^7^, 1*·*10*^−^*^6^, respectively). Once the role of these regulatory subunits in protein-phosphatase functioning in neurons is made clear, their contribution to SCZ-associated alterations of plasticity can be studied with our model as well. The predictions of our model could also be improved if detailed information on the relative expression of different isoforms of the plasticity regulating proteins was available.

In conclusion, our results show that the expression alterations observed in SCZ *post mortem*, especially those in anterior cingulate cortex, are likely to lead to impaired PKA-pathway-mediated potentiation in synapses containing GluR1 receptors. These results, validated by genomic and electrophysiological data from VEP modulation experiments, provide a possible genetic mechanism for plasticity impairments in SCZ and can form the basis of development of new pharmacological treatments.

## 4 Methods

### 4.1 Extension of the model of post-synaptic cortical plasticity with a description of neurogranin dynamics and model calibration

The model of [Måaki-Marttunen et al., 2020] described the three major intracellular signalling pathways mediating post-synaptic plasticity in the cortex, namely, the CaMKII, PKA, and PKC pathways. The model reproduced many types of neuromodulator-gated plasticity, including STDP, and due to the description of many involved proteins, it permits an evaluation of the contribution of many mental disorder-related genes to cortical plasticity. However, certain aspects of the model that are likely to be important for SCZ were unrealistic; in particular, the description of neurogranin was lacking detail and the basal levels of certain signalling molecules and receptors were underestimated. Here, we updated the model with adjustments required to improve the predictive power of the model in terms of applicability to SCZ.

#### 4.1.1 Neurogranin

Neurogranin is a small protein that is prevalent in mammalian brain and in the dendritic spines in particular [Watson et al., 1992], and it has been associated with the risk of SCZ [Stefansson et al., 2009, Ripke et al., 2014]. Neurogranin is phosphorylated by PKC and dephosphorylated by PP1, PP2A, and PP2B [Seki et al., 1995]. While the model of [Måaki-Marttunen et al., 2020] described neurogranin and its CaM buffering, it did not describe the phosphorylation of neurogranin by PKC and its consequences on CaM-affinity — a mechanism that regulates the type and magnitude of plasticity in the post-synaptic spine [Krucker et al., 2002, Huang et al., 2004]. Here, we extended the model to describe the PKC phosphorylation of neurogranin and its effects on CaM binding using in vitro biochemical data [Sheu et al., 1995, Seki et al., 1995].

First, we updated the reactions between CaM and neurogranin according to the current understanding of the interactions between these species. All forms of CaM (apo-CaM and CaM bound with 2, 3, or 4 Ca^2+^ ions) were allowed to bind to non-phosphorylated neurogranin, but the binding rate to Ca^2+^-bound CaM was 14 times slower and unbinding rate 4 times faster than to non-Ca^2+^-bound CaM, as in [Kubota et al., 2008]. We then adjusted the reaction rates of adding and removing of Ca^2+^ ions from CaM-bound neurogranin to keep the thermodynamical equilibrium. The phosphorylated neurogranin was not allowed to bind to any form of CaM, similar to [Kubota et al., 2008]. The reactions between neurogranin, CaM and Ca^2+^ are illustrated in Fig. 6A.

**Figure 6:**
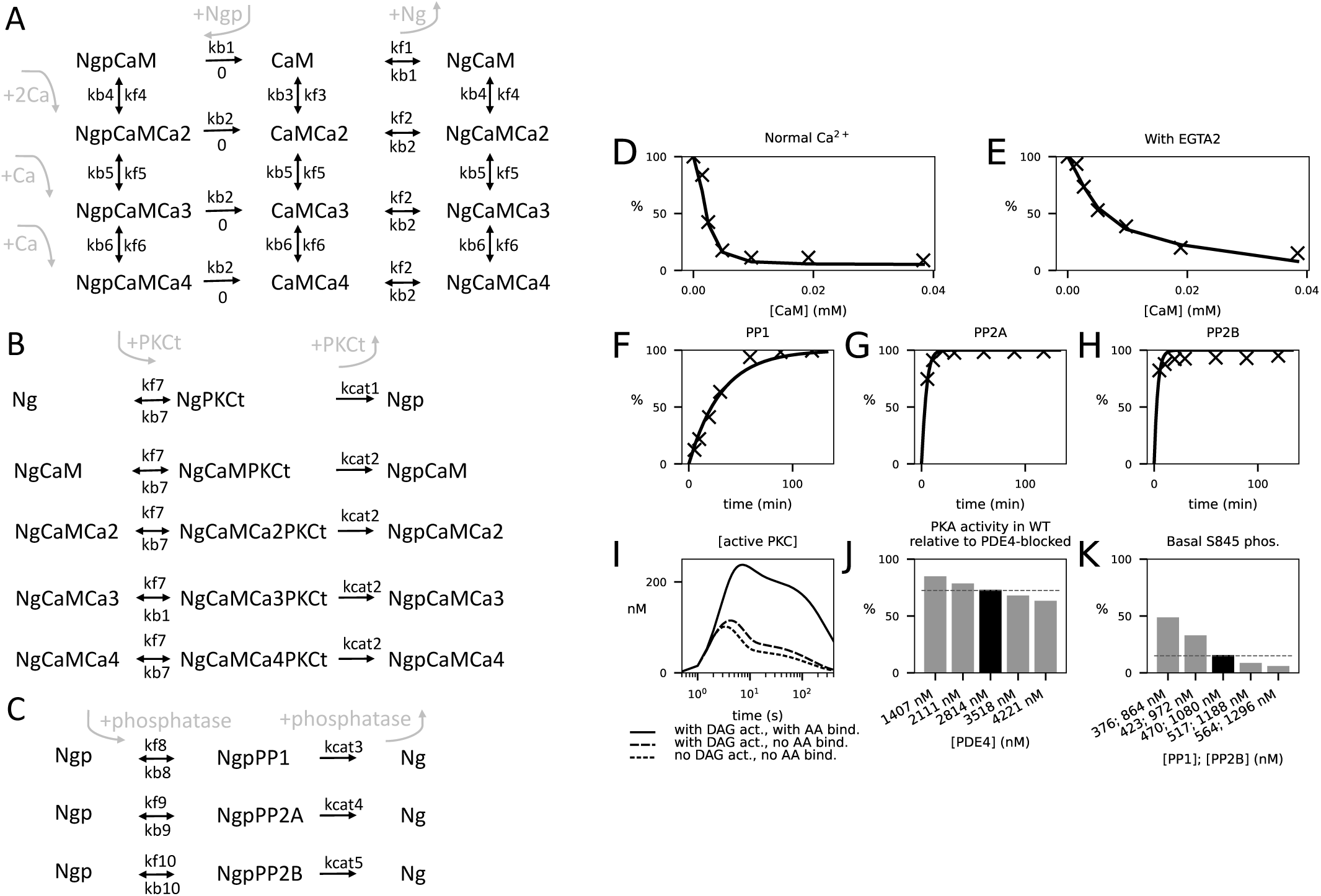
Extension of the model. **A**: Binding and unbinding reactions between CaM and phosphory-lated and non-phosphorylated neurogranin.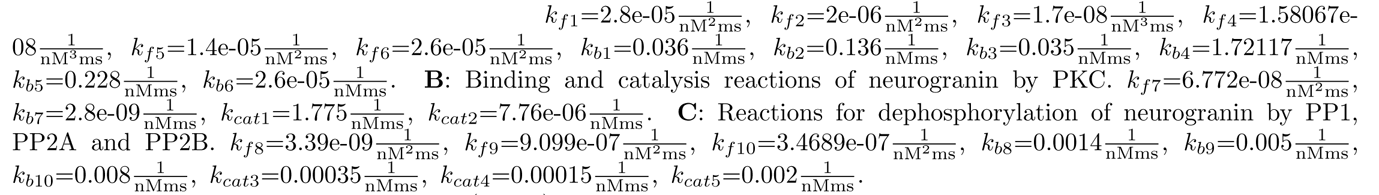, **D–E**: The model reproduces experimentaldata on phosphorylation of neurogranin (y-axis) under different concentrations of CaM (x-axis) in the absence (D) and presence (E) of EGTA2 (modelled as 90% less Ca^2+^). Solid curves represent model predictions, markers represent the data of [Sheu et al., 1995]. **F–H**: The model reproduces experimental time-course data on the dephosphorylation of neurogranin (y-axis) by PP1 (F), PP2A (G), and PP2B (H). Solid curves represent model predictions, markers represent the data of [Seki et al., 1995]. **I**: Effects of DAGK and AA activity on PKC activation. The solid line shows the time course of the concentration of active PKC with normal DAGK and AA activity and normal Gq-pathway neuromodulation. The dashed curves show the active PKC concentration when AA binding to DAG-bound PKC was disallowed and where the Gq-pathway neuromodulators to activate DAG were present (long dashes) or absent (short dashes). In these simulations, neurogranin concentration was decreased to 10% to highlight the PKC activation dynamics in a environment with reduced substrate load. **J**: Effects of PDE4 concentration on the ratio of amplitudes of PKA activation in the presence and absence of PDE4. The dashed line shows the experimentally observed value in prefrontal cortex [Castro et al., 2013]. **K**: Effects of protein phosphatase concentration on the basal level of S845-phosphorylated GluR1. The dashed line shows experimentally observed value [Oh et al., 2005].

Next, we fitted the rates of the reactions for phosphorylation of neurogranin by PKC. We adopted the catalysis rate *k_cat_* = 1.8/ms for NgPKC *→* Ngp + PKC as measured in [Sheu et al., 1995]. We varied the backward rate *k_b_* (of NgPKC *→* Ng + PKC) and controlled the forward rate (of Ng + PKC *→* NgPKC) as *k_f_* = *<u>^kb^</u>*<u>^+*k*^</u>*<u>^cat^</u>*, where *K_M_* = 26.2 mM was measured in [Sheu et al., 1995]. However, none of the models within this 1-dimensional parameter space fitted well to the data reported in [Sheu et al., 1995] under multiple CaM concentrations. Therefore, we allowed the CaM-bound neurogranin to be catalysed by PKC at a different rate than the non-CaM-bound neurogranin. We searched for the optimal model by a multi-objective optimisation algorithm (NSGA-II) by introducing an error function that quantified the difference between measured and predicted phosphorylation level both in absence and presence of a Ca^2+^ buffer (EGTA2) and running the optimizer for 25 generations. We obtained a good fit to experimental data across the CaM concentrations used in [Sheu et al., 1995] (Fig. 6D-E). The reactions between neurogranin and active PKC are illustrated in Fig. 6B.

To allow neurogranin activity to return to baseline level, we introduced the reactions for dephosphorylation of neurogranin by phosphatases PP1, PP2A and PP2B. We fitted the rates of Ngp binding with each of the three phosphatases to data from corresponding biochemical experiments [Seki et al., 1995], while we adopted the backward and catalysis rates from other reactions of the same phosphatase. The obtained dephosphorylation dynamics were in good agreement with experimental data (Fig. 6F–H). The reactions between neurogranin and the phosphatases are illustrated in Fig. 6C.

#### 4.1.2 Extended PKC activation scheme

The model of [Måaki-Marttunen et al., 2020] described the short-and long-term activation of PKC but lacked in the details on how they are dependent on Ca^2+^, DAG and AA. Experimental data show that transient PKC activation (translocation to the membrane) can be achieved by Ca^2+^ alone, and when accompanied by DAG activity, the PKC activity is sustained for a period of tens of seconds [Oancea and Meyer, 1998, Mogami et al., 2003]. Furthermore, when Ca^2+^ activity is accompanied by both DAG and AA, PKC activation lasts for minutes [Shirai et al., 1998]. We adopted the PKC activation model of [Gallimore et al., 2018] designed for reproduction of the data of [Oancea and Meyer, 1998, Mogami et al., 2003]. We removed the reactions adding a second Ca^2+^ ion to the Ca^2+^-bound PKC due to their small effect on the model dynamics. This simplification reduced the active forms of PKC to four from eight, and since active PKC interacts with many proteins in our model, this reduction led to omission of 112 reactions in total and thus significantly lessened the computational load of the model. Furthermore, we estimated the decay constant of the PKC activity from Figure 7C of [Shirai et al., 1998], and fitted the backward reaction rates of reaction PKCtDAGCa + AA *↔* PKCtAADAGCa (accompanied by a similar change in backward reaction rate of PKCtAACa + DAG *↔* PKCtAADAGCa to keep thermodynamic equilibrium) to these data. The model qualitatively reproduced the experimentally observed effects of DAG and AA activation on the duration of PKC activity (Fig. 6I): the PKC activity was short-lived (time above half-maximum *<* 10 s) when unaccompanied by DAG or AA activation [Oancea and Meyer, 1998], mildly longer (*≈*30 s) when accompanied by DAG activation [Oancea and Meyer, 1998], and significantly prolonged (*>* 3 min) when accompanied by both DAG activation and AA-binding of PKC [Shirai et al., 1998].

#### 4.1.3 Basal levels of Ca^2+^, S845-phosphorylated GluR1, and membrane-insertion of glutamate receptors

We further adjusted the model of [Måaki-Marttunen et al., 2020] to reproduce realistic basal levels of second messengers and phosphorylation of glutamate receptors. Recent models have used various concentrations for PMCA and NCX pumps (NCX concentration varied from 62 nM to 540 µM and PMCA concentration from 210 nM to 22 µM [Gallimore et al., 2018, Blackwell et al., 2018, Måaki-Marttunen et al., 2020]): here, we set the concentration of both to 6000 nM. Likewise, concentrations of adenylyl cyclases vary between 20 nM [Bhalla and Iyengar, 1999] and 6000 nM [Jedrzejewska-Szmek et al., 2017]: here, we set the concentration of AC1 (Gs-activated) to 200 nM. We also adjusted the concentration AC8 (non-Gs-activated) to a smaller value (100 nM) following the single-cell RNA expression data which suggests significantly smaller expression of ADCY8 in excitatory neurons than ADCY1 (Human Protein Atlas). We adjusted the leak Ca^2+^ channel concentration ([Leak] = 5130 nM) to obtain a basal Ca^2+^ concentration [Ca^2+^] = 50 nM. For the phosphodiesterase PDE4, we used a faster hydrolysis rate of cAMP from [Laliberté et al., 2002]. Based on a computational model [Ohadi et al., 2019], we included the PKA-mediated phosphorylation of PDE1, where the phosphorylated PDE1 had a 20 times smaller affinity to CaM than the non-phosphorylated PDE1 [Sharma and Wang, 1985], and used *K_m_*and *V*_max_ as measured in [Sharma and Wang, 1985]. We then fitted the PDE4-to-PDE1 concentration ratio to reproduce the effects of PDE4 blockade on the amplitude of PKA activation of [Castro et al., 2013] (Fig. 6J). As for basal levels of glutamate receptor phosphorylation and membrane insertion, we adjusted the PP1 and PP2B concentration (scaled down in proportion from the values used in [Måaki-Marttunen et al., 2020]) to get a basal level of 15% S845 phosphorylation, as observed in experiments [Oh et al., 2005] (Fig. 6K). Furthermore, we adopted the binding rate of activated PKC to GluR2 from the corresponding reaction between protein kinase and AMPAR subunit in [Jedrzejewska-Szmek et al., 2017] and used the membrane-insertion rate of non-phosphorylated GluR2 as given in [Gallimore et al., 2018]: this gave a good agreement with experimental data on the basal fraction of membrane-inserted GluR2s (42% predicted by the model, 45% measured in [Ashby et al., 2004]) and the magnitude of GluR2-endocytosis-mediated LTD [Chung et al., 2003].

### 4.2 Simulation framework

We modelled the spine (volume 0.5 fl) as a well-mixed, single compartment. The synapse was simulated using NEURON’s reaction-diffusion (RxD) extension [McDougal et al., 2013]. In most simulations of the biochemical model, we modelled the Ca^2+^ inputs to the spine as square-pulse currents of duration 3 ms and amplitude 100 Ca^2+^ ions/ms, unless otherwise stated. Each Ca^2+^ pulse was accompanied by a 3-ms pulse of *β*-adrenergic ligand, glutamate, and acetylcholine, unless otherwise stated.

The outputs of the biochemically detailed model were the time series of the concentrations of all molecular species, the membrane-inserted GluR subunits in particular. We used the statistical model for numbers of AMPAR tetramers at the membrane [Måaki-Marttunen et al., 2020] and the total synaptic conductance based on the predicted numbers of GluR subunits at the membrane and single-channel conductance of different types of tetramers [Oh and Derkach, 2005]. Namely, we determined the expected numbers of different types of tetramers as

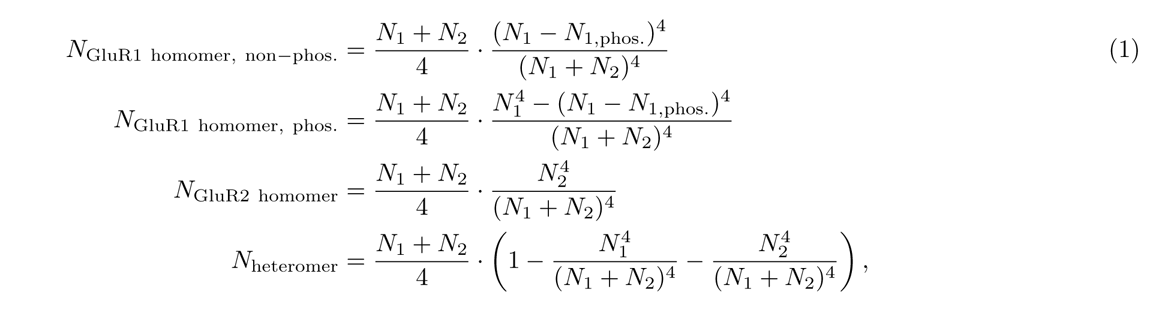

where *N*_1_ and *N*_2_ are the numbers of GluR1 and GluR2 subunits, respectively, at the membrane, and *N*_1,phos._ is the number of S831-phosphorylated GluR1 subunits at the membrane [Måaki-Marttunen et al., 2020]. The total synaptic conductance was determined as

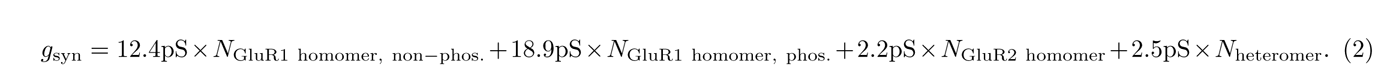

All simulations of the biochemically detailed model were first run for 6 hours without stimulation to reach a steady state. The relative synaptic conductance was quantified as the post-stimulus synaptic conductance of Eq. 2 divided by the synaptic conductance immediately before the start of the stimulation.

In certain simulations, we modelled realistic Ca^2+^ inputs to post-synaptic spines of L23PCs instead of the square-pulse Ca^2+^ inputs. To do this, we separately simulated a biophysically detailed multicompartmental model from [Markram et al., 2015] (L23 PC cADpyr229 1) to assess the amplitudes and time courses of the NMDAR-mediated Ca^2+^ inputs (see Supplementary material for details).

The model reactions and initial concentrations are listed in Tables S1 and S2, respectively. The NEURON/Python implementation of our model is publicly available in modelDB (accession number 267741).

### 4.3 Identification of the genes of interest and modelling their effects

In our previous work on neuronal effects of SCZ-associated genes, we identified the risk SNPs in genes encoding ion channels expressed in neocortical pyramidal cells and narrowed down the selection to the genes encoding channels described in our multicompartmental, Hodgkin-Huxley-based models [Måaki-Marttunen et al., 2016, Måaki-Marttunen et al., 2017, Måaki-Marttunen et al., 2019b]. Here, we repeated this procedure for genes relevant for synaptic plasticity in the cortex. We first listed all genes in the gene families encoding the proteins described in our model of neocortical plasticity [Måaki-Marttunen et al., 2020], complemented with SCZ risk genes associated with fast or slow neurotransmission [Devor et al., 2017] and the sister genes in their gene families (Table S4). We then sought for high-risk variants within these genes using the data of [Trubetskoy et al., 2022] (namely, from the “primary” meta-analysis of autosomal SNPs). We considered each gene where the minimal p-value for a single SNP was smaller than 5 *·* 10*^−^*^6^ as a gene of interest (Table 1A1).

A vast majority of the GWAS-identified SCZ-associated SNPs are non-coding variants, meaning that the risk variant results in a protein with identical structure to the non-risk protein [Harrison, 2015]. However, it is not certain that the risk is mediated by the altered expression of the gene within which, or closest to which, the SNP lies (see Discussion). The GWAS-identified SNPs may indeed affect expression of other genes through transcriptional regulation mechanisms (see, e.g., [Billingsley et al., 2018, McKinney et al., 2022]). We thus complemented the set of genes identified by GWAS (Table 1A1) by sets of genes that were differentially expressed in SCZ compared to HC. To do this, we analysed the CommonMind data sets of post-mortem RNA expression in PFC and ACC. We normalized the expression data using the DESeq2 method [Love et al., 2014]. We employed an imputation tool, CIBERSORTx [Steen et al., 2020], to obtain estimates of neuronal expression in these brain regions instead of the bulk expression using a reference single-cell RNA dataset [Zhang et al., 2016]. The imputation did not yield data for all genes due to low predicted expression or noise filtration [Steen et al., 2020] — we only considered the genes that were successfully imputed. We made a linear model for (imputed) expression of each gene of interest (Table S4) where we regressed against diagnosis (1=SCZ, 0=HC), age, sex (only individuals with XX and XY genotypes were included), and post-mortem interval (PMI). We considered each gene where the p-value of the regression was smaller than 5 *·* 10*^−^*^4^ as a hit — these genes are listed in Table 1A2 (PFC) and 1A3 (ACC). We used the exponentiated *β* coefficients for the diagnosis as a measure of *how much more* (exp(*β*) *>* 1.0) *or less* (exp(*β*) *<* 1.0) *expression of the gene there is to be expected in a subject with SCZ compared to a HC of the same age, sex, and PMI*. We used these data to inform our model on the direction and magnitude of the parameter change (Table 1, columns 3–5). In this work, we restricted our analysis on genes that showed a “medium” or “high” level of protein expression in neuronal cells of cerebral cortex *or* had a high specificity for expression in excitatory neurons according to the single-cell RNA sequencing data of Human Protein Atlas.

To allow extended analysis of the effects SCZ-associated variants on STDP, we repeated the analysis of Table 1A1–A3 for genes directly affecting the ion-channel currents described in the L23PC model (Table 1B1–B3) — see Supplementary material for details. When there were multiple SCZ hit genes affecting a single model parameter (e.g., DGKI and DGKZ in the ACC data, see Table 1A), we used the average of the expression-level factors to obtain a single factor.

For Figures 2H–K, 3L–O, and 4L–O, we performed subject-wise simulations to allow testing the statistical significance of the results. In these simulations, we altered the same parameters as shown in Table 1, but the initial concentration was set separately for each of the N=478 (ACC; 251 controls and 227 SCZ patients) or N=426 (PFC; 215 controls and 211 SCZ patients) samples. Namely, the initial concentration of protein *i* in subject *j* was multiplied by *x_i,j_/x_i,_*_HC_ where *x_i,j_* is the expression level of the protein and *x_i,_*_HC_ is the average expression level of the protein in the HC group.

### 4.4 Analysis of the phenotype-genotype data

To conduct the phenotype-genotype analyses, we used electrophysiological and genomic data from the Thematically Organised Psychosis (TOP) Study [Engh et al., 2010]. Five different gene set-based PRSs were calculated using beta coefficients and corresponding p-values from SCZ GWAS summary statistics [Trubetskoy et al., 2022] with TOP samples excluded. The largest set of genes included all genes from Table S4, namely, genes associated with plasticity and genes encoding ion channels or their subunits. The other sets were subsets of this set, namely, the set of plasticity-associated genes only, the set of ion channel-encoding genes only, the set of plasticity-associated genes that primarily coupled with the PKA pathway, and the set of plasticity-associated genes that primarily coupled with the PKC pathway (Table S4). For each of these sets, unless otherwise stated we included SNPs with SCZ-risk p-value *<* 10*^−^*^5^ and minor allele frequency *>* 0.01. No SNP pruning was performed.

The data consisted of N=286 HCs (114 male, 172 female), aged 49.9*±*16.9 years, that were not first-degree relatives to psychiatric patients. The VEP indices (C1, P1, and N1b amplitudes both at baseline and 2–4 min after the modulation block) were measured and determined from the Oz electrode as in [Valstad et al., 2020, Valstad et al., 2021]. The modulation of VEP was determined as the post-modulation C1, P1 or N1b amplitude subtracted from the baseline amplitude. A linear model was fit to the VEP modulation against one of the PRSs, age, and sex, and the hypothesis of association between the VEP modulation and the PRS (t-test, p-value threshold 0.05) was tested.

## Acknowledgements

Funding: Academy of Finland (330776, 336376, 318879), University of Oslo Convergence Environment (4MENT), and ERA-NET NEURON project SYNSCHIZ (Research Council of Norway, grant number 283798). **CommondMind**: Data were generated as part of the CommonMind Consortium supported by funding from Takeda Pharmaceuticals Company Limited, F. Hoffmann-La Roche Ltd and NIH grants R01MH085542, R01MH093725, P50MH066392, P50MH080405, R01MH097276, RO1-MH-075916, P50M096891, P50MH084053S1, R37MH057881, AG02219, AG05138, MH06692, R01MH110921, R01MH109677, R01MH109897, U01MH103392, and contract HHSN271201300031C through IRP NIMH. Brain tissue for the study was obtained from the following brain bank collections: the Mount Sinai NIH Brain and Tissue Repository, the University of Pennsylvania Alzheimer’s Disease Core Center, the University of Pittsburgh NeuroBioBank and Brain and Tissue Repositories, and the NIMH Human Brain Collection Core. CMC Leadership: Panos Roussos, Joseph Buxbaum, Andrew Chess, Schahram Akbarian, Vahram Haroutunian (Icahn School of Medicine at Mount Sinai), Bernie Devlin, David Lewis (University of Pittsburgh), Raquel Gur, Chang-Gyu Hahn (University of Pennsylvania), Enrico Domenici (University of Trento), Mette A. Peters, Solveig Sieberts (Sage Bionetworks), Thomas Lehner, Stefano Marenco, Barbara K. Lipska (NIMH).

