## Supplementary material for "Genetic mechanisms for impaired synaptic plasticity in schizophrenia revealed by computational modelling"

June 14, 2023

### Model reactions and initial conditions

Table S1: (Next page) Model reactions and reaction rates. Some of the reactions are grouped such that the variables $\mathbf{X}_i$, $\mathbf{Y}_i$ and/or $\mathbf{Z}_i$ represent the name or part of the name of the reactants or products of the reaction, given that they had the same reaction rates. See the bottom of the page for the declaration of the variables $\mathbf{X}_i$, $\mathbf{Y}_i$ and/or $\mathbf{Z}_i$. For example, Reaction 15 ($\ll$ $\mathbf{X}_1$ + PKAc $\rightleftharpoons$ PKAc$\mathbf{X}_1$ $\gg$) with $\mathbf{X}_1 \in$ {LR, pLR} is a short-hand notation for two reactions, $\ll$LR + PKAc $\rightleftharpoons$ PKAcLR$\gg$ and $\ll$pLR + PKAc $\rightleftharpoons$ PKAcpLR$\gg$; and Reaction 21 ($\ll$ $\mathbf{X}_3$ $\rightleftharpoons$ $\mathbf{Y}_3$ + Gibg$\gg$) with ($\mathbf{X}_3$, $\mathbf{Y}_3$) $\in$ { (ppppLRGibg, ppppLR), (ppppRGibg, ppppR) } is a short hand notation for two reactions, $\ll$ppppLRGibg $\rightleftharpoons$ ppppLR + Gibg$\gg$ and $\ll$ppppRGibg $\rightleftharpoons$ ppppR + Gibg$\gg$. For experiments where S845 phosphorylation of GluR1 by PKA was blocked, we introduced a spontaneous S845 phosphorylation GluR1$\mathbf{X}$ $\rightleftharpoons$ GluR1$\mathbf{X}$_$S$845, where $\mathbf{X}_1 \in$ {{}, _S831, _memb, _memb_S831} and the forward rate was fit to achieve a desired level of S845 (namely, PKA inactivation experiments of Fig. S1D where the rate was fit along other model parameters and the experiments of Fig. 1F,H where the rate was fit to make the GluR1 concentration at the membrane the same as without PKA inactivation). See ModelDB (http://modeldb.yale.edu/267741 — password "sczplast" needed during peer-review) for the full list of reactions in NeuroRD and NEURON formats as well as plain text.

| ID | Reaction | $k_f$ | $k_b$ | ID | Reaction | $k_f$ | $k_b$ |
|---|---|---|---|---|---|---|---|
| 1 | Ca + PMCA ⇌ PMCACa | 5e-05 | 0.007 | 40 | AC1CaMCa4ATP ⇌ cAMP + AC1CaMCa4 | 0.005684 | 0.0 |
| 2 | PMCACa ⇌ PMCA + CaOut | 0.0035 | 0.0 | 41 | AC8 + CaMCa4 ⇌ AC8CaMCa4 | 1.25e-06 | 0.001 |
| 3 | Ca + NCX ⇌ NCXCa | 1.68e-05 | 0.0112 | 42 | $\mathbf{X}_9$ + 2*Ca ⇌ $\mathbf{Z}_9$ | 1.7e-08 | 0.035 |
| 4 | NCXCa ⇌ NCX + CaOut | 0.0056 | 0.0 | 43 | $\mathbf{X}_{10}$ + Ca ⇌ $\mathbf{Z}_{10}$ | 1.4e-05 | 0.228 |
| 5 | CaOut + Leak ⇌ CaOutLeak | 1.5e-06 | 0.0011 | 44 | $\mathbf{X}_{11}$ + Ca ⇌ $\mathbf{Z}_{11}$ | 2.6e-05 | 0.064 |
| 6 | CaOutLeak ⇌ Ca + Leak | 0.0011 | 0.0 | 45 | Ng$\mathbf{X}_{12}$ + 2*Ca ⇌ Ng$\mathbf{Z}_{12}$ | 1.58067e-08 | 1.72117 |
| 7 | Ca + Calbin ⇌ CalbinC | 2.8e-05 | 0.0196 | 46 | CaM + Ng ⇌ NgCaM | 2.8e-05 | 0.036 |
| 8 | L ⇌ LOut | 0.0005 | 0.0 | 47 | CaM + Ngp ⇌ NgpCaM | 0.0 | 0.036 |
| 9 | L + R ⇌ LR | 5.555e-06 | 0.005 | 48 | $\mathbf{X}_{13}$ + Ng ⇌ Ng$\mathbf{X}_{13}$ | 2e-06 | 0.136 |
| 10 | LR + Gs ⇌ LRGs | 6e-07 | 1e-06 | 49 | $\mathbf{X}_{14}$ + Ngp ⇌ Ngp$\mathbf{X}_{14}$ | 0.0 | 0.136 |
| 11 | Gs + R ⇌ GsR | 4e-08 | 3e-07 | 50 | CaM$\mathbf{X}_{15}$ + PP2B ⇌ PP2BCaM$\mathbf{X}_{15}$ | 4.6e-07 | 1.2e-06 |
| 12 | GsR + L ⇌ LRGs | 2.5e-06 | 0.0005 | 51 | CaMCa3 + PP2B ⇌ PP2BCaMCa3 | 4.6e-06 | 1.2e-06 |
| 13 | LRGs ⇌ LRGsbg + GsaGTP | 0.02 | 0.0 | 52 | CaMCa4 + PP2B ⇌ PP2BCaMCa4 | 4.6e-05 | 1.2e-06 |
| 14 | LRGsbg ⇌ LR + Gsbg | 0.08 | 0.0 | 53 | PP2BCaMCa2 + Ca ⇌ PP2BCaMCa3 | 1.4e-05 | 0.0228 |
| 15 | $\mathbf{X}_1$ + PKAc ⇌ PKAc$\mathbf{X}_1$ | 8e-07 | 0.00448 | 54 | PP2BCaMCa3 + Ca ⇌ PP2BCaMCa4 | 2.6e-05 | 0.0064 |
| 16 | $\mathbf{X}_2$ ⇌ $\mathbf{Y}_2$ + PKAc | 0.001 | 0.0 | 55 | CaMCa4 + CK ⇌ CKCaMCa4 | 1e-05 | 0.003 |
| 17 | ppLR + PKAc ⇌ PKAcppLR | 1.712e-05 | 0.00448 | 56 | 2*CKCaMCa4 ⇌ Complex | 1e-07 | 0.01 |
| 18 | pppLR + PKAc ⇌ PKAcpppLR | 0.001712 | 0.00448 | 57 | CKpCaMCa4 + CKCaMCa4 ⇌ pComplex | 1e-07 | 0.01 |
| 19 | ppppLR + Gi ⇌ ppppLRGi | 0.00015 | 0.00025 | 58 | $\mathbf{X}_{16}$CaMCa4 + Complex | | |
| 20 | ppppLRGi ⇌ ppppLRGibg + GiaGTP | 0.000125 | 0.0 | | ⇌ $\mathbf{X}_{16}$CaMCa4 + pComplex | 1e-07 | 0.0 |
| 21 | $\mathbf{X}_3$ ⇌ $\mathbf{Y}_3$ + Gibg | 0.001 | 0.0 | 59 | 2*Complex ⇌ Complex + pComplex | 1e-05 | 0.0 |
| 22 | p$\mathbf{X}_4$ ⇌ $\mathbf{X}_4$ | 2.5e-06 | 0.0 | 60 | Complex + pComplex ⇌ 2*pComplex | 3e-05 | 0.0 |
| 23 | R + PKAc ⇌ PKAcR | 4e-08 | 0.00448 | 61 | CKpCaMCa4 ⇌ CaMCa4 + CKp | 8e-07 | 1e-05 |
| 24 | pR + PKAc ⇌ PKAcpR | 4e-07 | 0.00448 | 62 | CKp$\mathbf{X}_{17}$ + PP1 ⇌ CKp$\mathbf{Z}_{17}$ | 4e-09 | 0.00034 |
| 25 | ppR + PKAc ⇌ PKAcppR | 4e-06 | 0.00448 | 63 | CKp$\mathbf{X}_{18}$ ⇌ PP1 + $\mathbf{Z}_{18}$ | 8.6e-05 | 0.0 |
| 26 | pppR + PKAc ⇌ PKAcpppR | 0.0004 | 0.00448 | 64 | PKA + 4*cAMP ⇌ PKAcAMP4 | 1.6e-15 | 6e-05 |
| 27 | ppppR + Gi ⇌ ppppRGi | 7.5e-05 | 0.000125 | 65 | Epac1 + cAMP ⇌ Epac1cAMP | 3.1e-08 | 6.51e-05 |
| 28 | ppppRGi ⇌ ppppRGibg + GiaGTP | 6.25e-05 | 0.0 | 66 | I1 + PKAc ⇌ I1PKAc | 1.4e-06 | 0.0056 |
| 29 | GsaGTP ⇌ GsaGDP | 0.01 | 0.0 | 67 | I1PKAc ⇌ Ip35 + PKAc | 0.0014 | 0.0 |
| 30 | GsaGDP + Gsbg ⇌ Gs | 0.1 | 0.0 | 68 | Ip35 + PP1 ⇌ Ip35PP1 | 1e-06 | 1.1e-06 |
| 31 | GiaGTP ⇌ GiaGDP | 0.000125 | 0.0 | 69 | Ip35$\mathbf{X}_{19}$ + PP2BCaMCa4 ⇌ Ip35PP$\mathbf{Z}_{19}$ | 9.625e-05 | 0.33 |
| 32 | GiaGDP + Gibg ⇌ Gi | 0.00125 | 0.0 | 70 | Ip35PP$\mathbf{X}_{20}$ ⇌ I1 + PP$\mathbf{X}_{20}$ | 0.055 | 0.0 |
| 33 | GsaGTP + AC1 ⇌ AC1GsaGTP | 3.85e-05 | 0.01 | 71 | PP1PP2BCaMCa4 ⇌ PP1 + PP2BCaMCa4 | 0.0015 | 0.0 |
| 34 | AC1$\mathbf{X}_5$ + CaMCa4 ⇌ AC1$\mathbf{Z}_5$ | 6e-06 | 0.0009 | 72 | GluR1$\mathbf{X}_{21}$ + PKAc ⇌ GluR1$\mathbf{Z}_{21}$ | 4e-06 | 0.024 |
| 35 | $\mathbf{X}_6$CaMCa4 + ATP ⇌ $\mathbf{X}_6$CaMCa4ATP | 1e-05 | 0.07 | 73 | GluR1$\mathbf{X}_{22}$ ⇌ GluR1$\mathbf{Y}_{22}$ + PKAc | 0.006 | 0 |
| 36 | AC1GsaGTPCaMCa4ATP | | | 74 | GluR1$\mathbf{X}_{23}$ + $\mathbf{Y}_{23}$ ⇌ GluR1$\mathbf{Z}_{23}$ | 2.224e-08 | 0.0016 |
| | ⇌ cAMP + AC1GsaGTPCaMCa4 | 0.05684 | 0.0 | 75 | GluR1$\mathbf{X}_{24}$ ⇌ GluR1$\mathbf{Y}_{24}$ + $\mathbf{Z}_{24}$ | 0.0004 | 0 |
| 37 | AC1$\mathbf{X}_7$ + $\mathbf{Y}_7$ ⇌ AC1$\mathbf{Z}_7$ | 6.25e-05 | 0.01 | 76 | GluR1$\mathbf{X}_{25}$ + $\mathbf{Y}_{25}$ ⇌ GluR1$\mathbf{Z}_{25}$ | 2.78e-08 | 0.002 |
| 38 | $\mathbf{X}_8$CaMCa4ATP ⇌ cAMP + $\mathbf{X}_8$CaMCa4 | 0.002842 | 0.0 | 77 | GluR1$\mathbf{X}_{26}$ ⇌ GluR1$\mathbf{Y}_{26}$ + $\mathbf{Z}_{26}$ | 0.0005 | 0 |
| 39 | AC1GiaGTPCaMCa4ATP | | | 78 | GluR1$\mathbf{X}_{27}$ + PP1 ⇌ GluR1$\mathbf{Z}_{27}$ | 3e-08 | 0.00068 |
| | ⇌ cAMP + AC1GiaGTPCaMCa4 | 0.0005684 | 0.0 | 79 | GluR1$\mathbf{X}_{28}$ ⇌ GluR1$\mathbf{Y}_{28}$ + PP1 | 0.00017 | 0 |

| ID | Parameter names |
|---|---|
| 15 | $\mathbf{X}_1$ ∈ {LR, pLR} |
| 16 | ($\mathbf{X}_2$, $\mathbf{Y}_2$) ∈ { (PKAcLR, pLR), (PKAcpLR, ppLR), (PKAcppLR, pppLR), (PKAcpppLR, ppppLR), (PKAcR, pR), (PKAcpR, ppR), (PKAcppR, pppR), (PKAcpppR, ppppR), (PDE1A_PKAc, PDE1Ap) } |
| 21 | ($\mathbf{X}_3$, $\mathbf{Y}_3$) ∈ { (ppppLRGibg, ppppLR), (ppppRGibg, ppppR) } |
| 22 | $\mathbf{X}_4$ ∈ {LR, pLR, ppLR, pppLR, R, pR, ppR, pppR} |
| 34 | ($\mathbf{X}_5$, $\mathbf{Z}_5$) ∈ { (GsaGTP, GsaGTPCaMCa4), (GsaGTPGiaGTP, GsaGTPGiaGTPCaMCa4), ({}, CaMCa4) } |
| 35 | $\mathbf{X}_6$ ∈ {AC1GsaGTP, AC1GsaGTPGiaGTP, AC1GiaGTP, AC1, AC8} |
| 37 | ($\mathbf{X}_7$, $\mathbf{Y}_7$, $\mathbf{Z}_7$) ∈ { (GsaGTP, GiaGTP, GsaGTPGiaGTP), (CaMCa4, GiaGTP, GiaGTPCaMCa4), (GiaGTP, GsaGTP, GsaGTPGiaGTP) } |
| 38 | $\mathbf{X}_8$ ∈ {AC1GsaGTPGiaGTP, AC8} |
| 42 | ($\mathbf{X}_9$, $\mathbf{Z}_9$) ∈ { (CaM, CaMCa2), (PP2BCaM, PP2BCaMCa2) } |
| 43 | ($\mathbf{X}_{10}$, $\mathbf{Z}_{10}$) ∈ { (CaMCa2, CaMCa3), (NgCaMCa2, NgCaMCa3), (NgpCaMCa2, NgpCaMCa3) } |
| 44 | ($\mathbf{X}_{11}$, $\mathbf{Z}_{11}$) ∈ { (CaMCa3, CaMCa4), (NgCaMCa3, NgCaMCa4), (NgpCaMCa3, NgpCaMCa4) } |
| 45 | ($\mathbf{X}_{12}$, $\mathbf{Z}_{12}$) ∈ { (CaM, CaMCa2), (pCaM, pCaMCa2) } |
| 48 | $\mathbf{X}_{13}$ ∈ {CaMCa2, CaMCa3, CaMCa4} |
| 49 | $\mathbf{X}_{14}$ ∈ {CaMCa2, CaMCa3, CaMCa4} |
| 50 | $\mathbf{X}_{15}$ ∈ {{}, Ca2} |
| 58 | $\mathbf{X}_{16}$ ∈ {CKp, CK} |
| 62 | ($\mathbf{X}_{17}$, $\mathbf{Z}_{17}$) ∈ { ({}, PP1), (CaMCa4, CaMCa4PP1) } |
| 63 | ($\mathbf{X}_{18}$, $\mathbf{Z}_{18}$) ∈ { (PP1, CK), (CaMCa4PP1, CKCaMCa4) } |
| 69 | ($\mathbf{X}_{19}$, $\mathbf{Z}_{19}$) ∈ { ({}, 2BCaMCa4), (PP1, 1PP2BCaMCa4) } |
| 70 | $\mathbf{X}_{20}$ ∈ {2BCaMCa4, 1PP2BCaMCa4} |
| 72 | ($\mathbf{X}_{21}$, $\mathbf{Z}_{21}$) ∈ { ({}, _PKAc), (_S831, _S831_PKAc), (_memb, _memb_PKAc), (_memb_S831, _memb_S831_PKAc) } |
| 73 | ($\mathbf{X}_{22}$, $\mathbf{Y}_{22}$) ∈ { (_PKAc, _S845), (_S831_PKAc, _S845_S831), (_memb_PKAc, _memb_S845), (_memb_S831_PKAc, _memb_S845_S831) } |
| 74 | ($\mathbf{X}_{23}$, $\mathbf{Y}_{23}$, $\mathbf{Z}_{23}$) ∈ { ({}, CKCaMCa4, _CKCaMCa4), ({}, CKp, _CKp), (_S845, CKCaMCa4, _S845_CKCaMCa4), (_S845, CKp, _S845_CKp), (_memb, CKCaMCa4, _memb_CKCaMCa4), (_memb, CKp, _memb_CKp), (_memb_S845, CKCaMCa4, _memb_S845_CKCaMCa4), (_memb_S845, CKp, _memb_S845_CKp) } |
| 75 | ($\mathbf{X}_{24}$, $\mathbf{Y}_{24}$, $\mathbf{Z}_{24}$) ∈ { (_CKCaMCa4, _S831, CKCaMCa4), (_CKp, _S831, CKp), (_S845_CKCaMCa4, _S845_S831, CKCaMCa4), (_S845_CKp, _S845_S831, CKp), (_memb_CKCaMCa4, _memb_S831, CKCaMCa4), (_memb_CKp, _memb_S831, CKp), (_memb_S845_CKCaMCa4, _memb_S845_S831, CKCaMCa4), (_memb_S845_CKp, _memb_S845_S831, CKp) } |
| 76 | ($\mathbf{X}_{25}$, $\mathbf{Y}_{25}$, $\mathbf{Z}_{25}$) ∈ { ({}, CKpCaMCa4, _CKpCaMCa4), ({}, PKCtCa, _PKCtCa), ({}, PKCtAACa, _PKCtAACa), ({}, PKCtDAGCa, _PKCtDAGCa), ({}, PKCtAADAGCa, _PKCtAADAGCa), (_S845, CKpCaMCa4, _S845_CKpCaMCa4), (_S845, PKCtCa, _S845_PKCtCa), (_S845, PKCtAACa, _S845_PKCtAACa), (_S845, PKCtDAGCa, _S845_PKCtDAGCa), (_S845, PKCtAADAGCa, _S845_PKCtAADAGCa), (_memb, CKpCaMCa4, _memb_CKpCaMCa4), (_memb, PKCtCa, _memb_PKCtCa), (_memb, PKCtAACa, _memb_PKCtAACa), (_memb, PKCtDAGCa, _memb_PKCtDAGCa), (_memb, PKCtAADAGCa, _memb_PKCtAADAGCa), (_memb_S845, CKpCaMCa4, _memb_S845_CKpCaMCa4), (_memb_S845, PKCtCa, _memb_S845_PKCtCa), (_memb_S845, PKCtAACa, _memb_S845_PKCtAACa), (_memb_S845, PKCtDAGCa, _memb_S845_PKCtDAGCa), (_memb_S845, PKCtAADAGCa, _memb_S845_PKCtAADAGCa) } |
| 77 | ($\mathbf{X}_{26}$, $\mathbf{Y}_{26}$, $\mathbf{Z}_{26}$) ∈ { (_CKpCaMCa4, _S831, CKpCaMCa4), (_PKCtCa, _S831, PKCtCa), (_PKCtAACa, _S831, PKCtAACa), (_PKCtDAGCa, _S831, PKCtDAGCa), (_PKCtAADAGCa, _S831, PKCtAADAGCa), (_S845_CKpCaMCa4, _S845_S831, CKpCaMCa4), (_S845_PKCtCa, _S845_S831, PKCtCa), (_S845_PKCtAACa, _S845_S831, PKCtAACa), (_S845_PKCtDAGCa, _S845_S831, PKCtDAGCa), (_S845_PKCtAADAGCa, _S845_S831, PKCtAADAGCa), (_memb_CKpCaMCa4, _memb_S831, CKpCaMCa4), (_memb_PKCtCa, _memb_S831, PKCtCa), (_memb_PKCtAACa, _memb_S831, PKCtAACa), (_memb_PKCtDAGCa, _memb_S831, PKCtDAGCa), (_memb_PKCtAADAGCa, _memb_S831, PKCtAADAGCa), (_memb_S845_CKpCaMCa4, _memb_S845_S831, CKpCaMCa4), (_memb_S845_PKCtCa, _memb_S845_S831, PKCtCa), (_memb_S845_PKCtAACa, _memb_S845_S831, PKCtAACa), (_memb_S845_PKCtDAGCa, _memb_S845_S831, PKCtDAGCa), (_memb_S845_PKCtAADAGCa, _memb_S845_S831, PKCtAADAGCa) } |
| 78 | ($\mathbf{X}_{27}$, $\mathbf{Z}_{27}$) ∈ { (_S845, _S845_PP1), (_memb_S845, _memb_S845_PP1) } |
| 79 | ($\mathbf{X}_{28}$, $\mathbf{Y}_{28}$) ∈ { (_S845_PP1, {}), (_memb_S845_PP1, _memb) } |

| ID | Reaction | $k_f$ | $k_b$ |
|---|---|---|---|
| 80 | GluR1$\mathbf{X}_{29}$ + PP1 ⇌ GluR1$\mathbf{Z}_{29}$ | 3e-08 | 0.0014 |
| 81 | GluR1$\mathbf{X}_{30}$ ⇌ GluR1$\mathbf{Y}_{30}$ + PP1 | 0.00035 | 0 |
| 82 | GluR1$\mathbf{X}_{31}$ + PP2BCaMCa4 ⇌ GluR1$\mathbf{Z}_{31}$ | 2.01e-06 | 0.008 |
| 83 | $\mathbf{X}_{32}$ ⇌ $\mathbf{Y}_{32}$ + PP2BCaMCa4 | 0.002 | 0 |
| 84 | GluR1_memb$\mathbf{X}_{33}$ ⇌ GluR1$\mathbf{X}_{33}$ | 8e-07 | 0 |
| 85 | GluR1_S845$\mathbf{X}_{34}$ ⇌ GluR1_memb_S845$\mathbf{X}_{34}$ | 3.28e-05 | 8e-07 |
| 86 | PDE1A + CaMCa4 ⇌ PDE1ACaMCa4 | 0.0001 | 5e-05 |
| 87 | PDE1A$\mathbf{X}_{35}$ + cAMP ⇌ PDE1A$\mathbf{Z}_{35}$ | 2.37e-05 | 0.664 |
| 88 | PDE1A$\mathbf{X}_{36}$ ⇌ PDE1A$\mathbf{Y}_{36}$ + AMP | 0.166 | 0.0 |
| 89 | PDE1A + PKAc ⇌ PDE1A_PKAc | 1.19e-05 | 0.004 |
| 90 | PDE1Ap + CaMCa4 ⇌ PDE1ApCaMCa4 | 0.0001 | 0.001 |
| 91 | PDE1Ap ⇌ PDE1A | 0.00012 | 0.0 |
| 92 | AMP ⇌ ATP | 0.001 | 0.0 |
| 93 | PDE4 + cAMP ⇌ PDE4cAMP | 5e-08 | 0.00016 |
| 94 | PDE4cAMP ⇌ PDE4 + AMP | 4e-05 | 0.0 |
| 95 | $\mathbf{X}_{37}$ + $\mathbf{Y}_{37}$ ⇌ PKAc$\mathbf{Z}_{37}$ | 2.5e-07 | 8e-05 |
| 96 | PKAc$\mathbf{X}_{38}$ ⇌ PDE4p$\mathbf{Y}_{38}$ + PKAc | 2e-05 | 0.0 |
| 97 | PDE4p ⇌ PDE4 | 2e-05 | 0.0 |
| 98 | PDE4p + cAMP ⇌ PDE4pcAMP | 2e-07 | 0.00064 |
| 99 | PDE4pcAMP ⇌ PDE4p + AMP | 0.00016 | 0.0 |
| 100 | PKAcAMP4 ⇌ PKAr + 2*PKAc | 0.00024 | 2.55e-05 † |
| 101 | Ca + fixedbuffer ⇌ fixedbufferCa | 0.0004 | 20.0 |
| 102 | Glu ⇌ GluOut | 0.0005 | 0.0 |
| 103 | Ca + PLC ⇌ PLCCa | 4e-07 | 0.001 |
| 104 | GqaGTP + PLC ⇌ PLCGqaGTP | 7e-07 | 0.0007 |
| 105 | Ca + PLCGqaGTP ⇌ PLCCaGqaGTP | 8e-05 | 0.04 |
| 106 | GqaGTP + PLCCa ⇌ PLCCaGqaGTP | 0.0001 | 0.01 |
| 107 | PLCCa + Pip2 ⇌ PLCCaPip2 | 3e-08 | 0.01 |
| 108 | PLCCaPip2 ⇌ PLCCaDAG + Ip3 | 0.0003 | 0.0 |
| 109 | PLCCaDAG ⇌ PLCCa + DAG | 0.2 | 0.0 |
| 110 | PLCCaGqaGTP + Pip2 ⇌ PLCCaGqaGTPPip2 | 1.5e-05 | 0.075 |
| 111 | PLCCaGqaGTPPip2 ⇌ PLCCaGqaGTPDAG + Ip3 | 0.25 | 0.0 |
| 112 | PLCCaGqaGTPDAG ⇌ PLCCaGqaGTP + DAG | 1.0 | 0.0 |
| 113 | Ip3degrad + PIkinase ⇌ Ip3degPIk | 2e-06 | 0.001 |
| 114 | Ip3degPIk ⇌ PIkinase + Pip2 | 0.001 | 0.0 |
| 115 | PLC$\mathbf{X}_{39}$ ⇌ PLC$\mathbf{Y}_{39}$ + GqaGTP | 0.012 | 0.0 |
| 116 | GqaGTP ⇌ GqaGDP | 0.001 | 0.0 |
| 117 | GqaGDP ⇌ Gqabg | 0.01 | 0.0 |
| 118 | Ca + DGL ⇌ CaDGL | 0.000125 | 0.05 |
| 119 | DAG + CaDGL ⇌ DAGCaDGL | 5e-07 | 0.001 |
| 120 | DAGCaDGL ⇌ CaDGL + 2AG | 0.00025 | 0.0 |
| 121 | Ip3 ⇌ Ip3degrad | 0.01 | 0.0 |
| 122 | 2AG ⇌ 2AGdegrad | 0.005 | 0.0 |
| 123 | DAG + DAGK ⇌ DAGKdag | 7e-08 | 0.0008 |
| 124 | DAGKdag ⇌ PA + DAGK | 0.0002 | 0.0 |
| 125 | Ca + PKC ⇌ PKCCa | 1.33e-05 | 0.012 |
| 126 | $\mathbf{X}_{40}$ + AA ⇌ $\mathbf{Z}_{40}$ | 4e-06 | 0.001 |
| 127 | PKCCa ⇌ PKCtCa | 0.0 | 0.005 |
| 128 | PKCAACa ⇌ PKCtAACa | 0.003 | 0.005 |
| 129 | PKCtCa + DAG ⇌ PKCtDAGCa | 8e-07 | 0.001 |
| 130 | PKCtAACa + DAG ⇌ PKCtAADAGCa | 8e-07 | 5e-05 |
| 131 | PKCtDAGCa + AA ⇌ PKCtAADAGCa | 4e-06 | 5e-05 |
| 132 | Glu + MGluR ⇌ MGluR_Glu | 1.68e-08 | 0.0001 |
| 133 | MGluR_Glu ⇌ MGluR_Glu_desens | 6.25e-05 | 1e-06 |
| 134 | Gqabg + MGluR_Glu ⇌ MGluR_Gqabg_Glu | 9e-06 | 0.00136 |
| 135 | MGluR_Gqabg_Glu ⇌ GqaGTP + MGluR_Glu | 0.0015 | 0.0 |
| 136 | GluR2$\mathbf{X}_{41}$ + PKCt$\mathbf{Y}_{41}$ ⇌ GluR2$\mathbf{Z}_{41}$ | 4e-06 | 0.0008 |
| 137 | GluR2$\mathbf{X}_{42}$ ⇌ GluR2$\mathbf{Y}_{42}$ + PKCt$\mathbf{Z}_{42}$ | 0.0047 | 0 |
| 138 | GluR2$\mathbf{X}_{43}$ + PP2A ⇌ GluR2$\mathbf{Z}_{43}$ | 5e-07 | 0.005 |
| 139 | $\mathbf{X}_{44}$ ⇌ $\mathbf{Y}_{44}$ + PP2A | 0.00015 | 0 |
| 140 | GluR2$\mathbf{X}_{45}$ ⇌ GluR2_memb$\mathbf{X}_{45}$ | 0.00024545 | 0.0003 |
| 141 | GluR2_S880$\mathbf{X}_{46}$ ⇌ GluR2_memb_S880$\mathbf{X}_{46}$ | 0.0055 | 0.07 |
| 142 | ACh + M1R ⇌ AChM1R | 9.5e-08 | 0.0025 |
| 143 | Gqabg + AChM1R ⇌ AChM1RGq | 2.4e-05 | 0.00042 |
| 144 | Gqabg + M1R ⇌ M1RGq | 5.76e-07 | 0.00042 |
| 145 | ACh + M1RGq ⇌ AChM1RGq | 3.96e-06 | 0.0025 |
| 146 | AChM1RGq ⇌ GqaGTP + AChM1R | 0.0005 | 0.0 |
| 147 | ACh ⇌ | 0.0005 | 0 |
| 148 | Ca + PLA2 ⇌ CaPLA2 | 1.77e-05 | 0.154 |
| 149 | Ca + CaPLA2 ⇌ Ca2PLA2 | 1.63e-05 | 0.102 |
| 150 | Ca2PLA2 ⇌ Ca2PLA2act | 0.0015 | 0.008 |
| 151 | Ca2PLA2act + Pip2 ⇌ Ca2PLA2actPip2 | 2.2e-05 | 0.444 |
| 152 | Ca2PLA2actPip2 ⇌ Ca2PLA2act + AA | 0.111 | 0.0 |
| 153 | AA ⇌ Pip2 | 0.01 | 0.0 |
| 154 | Ng$\mathbf{X}_{47}$ + PKCt$\mathbf{Y}_{47}$ ⇌ Ng$\mathbf{Z}_{47}$ | 6.772e-08 | 2.8e-09 |
| 155 | NgPKCt$\mathbf{X}_{48}$ ⇌ Ngp + PKCt$\mathbf{X}_{48}$ | 1.775 | 0 |
| 156 | NgCaM$\mathbf{X}_{49}$ ⇌ NgpCaM$\mathbf{Y}_{49}$ + PKCt$\mathbf{Z}_{49}$ | 7.76e-06 | 0 |
| 157 | Ngp + PP1 ⇌ NgpPP1 | 3.39e-09 | 0.0014 |
| 158 | NgpPP1 ⇌ Ng + PP1 | 0.00035 | 0 |
| 159 | Ngp + PP2A ⇌ NgpPP2A | 9.099e-07 | 0.005 |
| 160 | Ngp + PP2BCaMCa4 ⇌ NgpPP2BCaMCa4 | 3.4689e-07 | 0.008 |

| ID | Parameter names |
|---|---|
| 80 | ($\mathbf{X}_{29}$, $\mathbf{Z}_{29}$) ∈ { (_S845_S831, _S845_S831_PP1), (_S831, _S831_PP1), (_memb_S845_S831, _memb_S845_S831_PP1), (_memb_S831, _memb_S831_PP1) } |
| 81 | ($\mathbf{X}_{30}$, $\mathbf{Y}_{30}$) ∈ { (_S845_S831_PP1, _S845), (_S845_S831_PP1, _S831), (_S831_PP1, {}), (_memb_S845_S831_PP1, _memb_S845), (_memb_S845_S831_PP1, _memb_S831), (_memb_S831_PP1, _memb) } |
| 82 | ($\mathbf{X}_{31}$, $\mathbf{Z}_{31}$) ∈ { (_S845, _S845_PP2B), (_S845_S831, _S845_S831_PP2B), (_memb_S845, _memb_S845_PP2B), (_memb_S845_S831, _memb_S845_S831_PP2B) } |
| 83 | ($\mathbf{X}_{32}$, $\mathbf{Y}_{32}$) ∈ { (GluR1_S845_PP2B, GluR1), (GluR1_S845_S831_PP2B, GluR1_S831), (GluR1_memb_S845_PP2B, GluR1_memb), (GluR1_memb_S845_S831_PP2B, GluR1_memb_S831), (NgpPP2BCaMCa4, Ng) } |
| 84 | $\mathbf{X}_{33}$ ∈ {{}, _PKAc, _CKCaMCa4, _CKpCaMCa4, _CKp, _PKCtCa, _PKCtAACa, _PKCtDAGCa, _PKCtAADAGCa, _S831, _S831_PKAc, _S831_PP1} |
| 85 | $\mathbf{X}_{34}$ ∈ {{}, _CKCaMCa4, _CKpCaMCa4, _CKp, _PKCtCa, _PKCtAACa, _PKCtDAGCa, _PKCtAADAGCa, _S831, _PP1, _S831_PP1, _PP2B, _S831_PP2B} |
| 87 | ($\mathbf{X}_{35}$, $\mathbf{Z}_{35}$) ∈ { (CaMCa4, CaMCa4cAMP), (pCaMCa4, pCaMCa4cAMP) } |
| 88 | ($\mathbf{X}_{36}$, $\mathbf{Y}_{36}$) ∈ { (CaMCa4cAMP, CaMCa4), (pCaMCa4cAMP, pCaMCa4) } |
| 95 | ($\mathbf{X}_{37}$, $\mathbf{Y}_{37}$, $\mathbf{Z}_{37}$) ∈ { (PKAc, PDE4, PDE4), (PDE4cAMP, PKAc, _PDE4_cAMP) } |
| 96 | ($\mathbf{X}_{38}$, $\mathbf{Y}_{38}$) ∈ { (PDE4, {}), (_PDE4_cAMP, cAMP) } |
| 115 | ($\mathbf{X}_{39}$, $\mathbf{Y}_{39}$) ∈ { (GqaGTP, {}), (CaGqaGTP, Ca) } |
| 126 | ($\mathbf{X}_{40}$, $\mathbf{Z}_{40}$) ∈ { (PKCCa, PKCAACa), (PKCtCa, PKCtAACa) } |
| 136 | ($\mathbf{X}_{41}$, $\mathbf{Y}_{41}$, $\mathbf{Z}_{41}$) ∈ { ({}, Ca, _PKCtCa), ({}, AACa, _PKCtAACa), ({}, DAGCa, _PKCtDAGCa), ({}, AADAGCa, _PKCtAADAGCa), (_memb, Ca, _memb_PKCtCa), (_memb, AACa, _memb_PKCtAACa), (_memb, DAGCa, _memb_PKCtDAGCa), (_memb, AADAGCa, _memb_PKCtAADAGCa) } |
| 137 | ($\mathbf{X}_{42}$, $\mathbf{Y}_{42}$, $\mathbf{Z}_{42}$) ∈ { (_PKCtCa, _S880, Ca), (_PKCtAACa, _S880, AACa), (_PKCtDAGCa, _S880, DAGCa), (_PKCtAADAGCa, _S880, AADAGCa), (_memb_PKCtCa, _memb_S880, Ca), (_memb_PKCtAACa, _memb_S880, AACa), (_memb_PKCtDAGCa, _memb_S880, DAGCa), (_memb_PKCtAADAGCa, _memb_S880, AADAGCa) } |
| 138 | ($\mathbf{X}_{43}$, $\mathbf{Z}_{43}$) ∈ { (_S880, _S880_PP2A), (_memb_S880, _memb_S880_PP2A) } |
| 139 | ($\mathbf{X}_{44}$, $\mathbf{Y}_{44}$) ∈ { (GluR2_S880_PP2A, GluR2), (GluR2_memb_S880_PP2A, GluR2_memb), (NgpPP2A, Ng) } |
| 140 | $\mathbf{X}_{45}$ ∈ {{}, _PKCtCa, _PKCtAACa, _PKCtDAGCa, _PKCtAADAGCa} |
| 141 | $\mathbf{X}_{46}$ ∈ {{}, _PP2A} |
| 154 | ($\mathbf{X}_{47}$, $\mathbf{Y}_{47}$, $\mathbf{Z}_{47}$) ∈ { ({}, Ca, PKCtCa), (CaM, Ca, CaMPKCtCa), (CaMCa2, Ca, CaMCa2PKCtCa), (CaMCa3, Ca, CaMCa3PKCtCa), (CaMCa4, Ca, CaMCa4PKCtCa), ({}, AACa, PKCtAACa), (CaM, AACa, CaMPKCtAACa), (CaMCa2, AACa, CaMCa2PKCtAACa), (CaMCa3, AACa, CaMCa3PKCtAACa), (CaMCa4, AACa, CaMCa4PKCtAACa), ({}, DAGCa, PKCtDAGCa), (CaM, DAGCa, CaMPKCtDAGCa), (CaMCa2, DAGCa, CaMCa2PKCtDAGCa), (CaMCa3, DAGCa, CaMCa3PKCtDAGCa), (CaMCa4, DAGCa, CaMCa4PKCtDAGCa), ({}, AADAGCa, PKCtAADAGCa), (CaM, AADAGCa, CaMPKCtAADAGCa), (CaMCa2, AADAGCa, CaMCa2PKCtAADAGCa), (CaMCa3, AADAGCa, CaMCa3PKCtAADAGCa), (CaMCa4, AADAGCa, CaMCa4PKCtAADAGCa) } |
| 155 | $\mathbf{X}_{48}$ ∈ {Ca, AACa, DAGCa, AADAGCa} |
| 156 | ($\mathbf{X}_{49}$, $\mathbf{Y}_{49}$, $\mathbf{Z}_{49}$) ∈ { (PKCtCa, {}, Ca), (Ca2PKCtCa, Ca2, Ca), (Ca3PKCtCa, Ca3, Ca), (Ca4PKCtCa, Ca4, Ca), (PKCtAACa, {}, AACa), (Ca2PKCtAACa, Ca2, AACa), (Ca3PKCtAACa, Ca3, AACa), (Ca4PKCtAACa, Ca4, AACa), (PKCtDAGCa, {}, DAGCa), (Ca2PKCtDAGCa, Ca2, DAGCa), (Ca3PKCtDAGCa, Ca3, DAGCa), (Ca4PKCtDAGCa, Ca4, DAGCa), (PKCtAADAGCa, {}, AADAGCa), (Ca2PKCtAADAGCa, Ca2, AADAGCa), (Ca3PKCtAADAGCa, Ca3, AADAGCa), (Ca4PKCtAADAGCa, Ca4, AADAGCa) } |

Table S2: Initial concentrations in the default model. All unnamed species had an initial concentration of 0 nM. The values used are the same as in [Mäki-Marttunen et al., 2020], except the ones adjusted, see Methods.

| Species | Concentration (nM) |
|---|---|
| CaOut | 1900000 |
| Leak | 5130 |
| Calbin | 150000 |
| CalbinC | 15000 |
| LOut | 2500000 |
| Epac1 | 500 |
| PMCA | 6000 |
| NCX | 6000 |
| L | 10 |
| R | 1600 |
| Gs | 13000 |
| Gi | 2600 |
| AC1 | 200 |
| ATP | 2000000 |
| AC8 | 100 |
| PDE1A | 12000 |

| Species | Concentration (nM) |
|---|---|
| AMP | 980 |
| Ng | 20000 |
| CaM | 60000 |
| PP2B | 987 |
| CK | 23000 |
| PKA | 6400 |
| I1 | 2200 |
| PP1 | 429 |
| GluR1 | 180 |
| GluR1_memb | 90 |
| PDE4 | 3039 |
| fixedbuffer | 500000 |
| MGluR | 800 |
| GluOut | 1000000 |
| Gqabg | 1400 |

| Species | Concentration (nM) |
|---|---|
| PLC | 250 |
| Pip2 | 24000 |
| PIkinase | 290 |
| Ip3degPIk | 400 |
| PKC | 15000 |
| DAG | 90 |
| DAGK | 300 |
| DGL | 1600 |
| CaDGL | 250 |
| DAGCaDGL | 90 |
| Ip3degrad | 600 |
| GluR2 | 14 |
| GluR2_memb | 256 |
| PP2A | 500 |
| M1R | 450 |
| PLA2 | 1000 |

### Modelling of post-synaptic $Ca^{2+}$ transients

To model the post-synaptic $Ca^{2+}$ transients in L23PCs in the STDP protocol, we simulated the single-cell model of L23PC as described in [Markram et al., 2015]. We inserted a dendritic spine (head diameter 0.85 μm, head length 0.85 μm, neck diameter 0.1 μm, neck length 0.5 μm) at a random location along the apical dendrite at distance 250–300 μm from the soma. We used the probabilistic AMPAR–NMDAR synapse model of [Hay and Segev, 2015] with $Mg^{2+}$ gating of NMDAR from [Spruston et al., 1995], a corrected pre-synaptic resource update rule [Mäki-Marttunen et al., 2018] and an AMPAR–NMDAR ratio of 1:1. We stimulated the spine head with synaptic stimuli every 1 s, which was coupled (interstimulus interval ranging from -200 to 200 ms) with a burst of 4 somatic stimuli. The AMPARs and NMDARs had both a maximal conductance of 150 pS and reversal potential 0 mV, rise time constants 0.3 and 2 ms, respectively, and decay time constants 3 and 65 ms, respectively [Markram et al., 2015]. The synapse obeyed a probabilistic short-term plasticity model with depression time constant $\tau_{\mathrm{dep}} = 650$ ms, facilitation time constant $\tau_{\mathrm{dep}} = 17$ ms, and resource use fraction $U_{\mathrm{se}} = 0.5$ [Markram et al., 2015]. TTX-sensitive $Na^{+}$ currents that amplify and prolong the post-synaptic currents have been observed in dendritic spines of neocortical pyramidal cells [Araya et al., 2007]. We thus introduced a persistent $Na^{+}$ current with maximal conductance 0.03 pS/cm$^2$ in the spine head. We repeated the coupled stimulus 100 times in each simulation, and calculated the numbers of $Ca^{2+}$ ions entering into the post-synaptic spine across time based on the synaptic currents from this model — assuming 0% of AMPAR-conducted current and 10% of NMDAR-conducted current was $Ca^{2+}$. We further averaged the $Ca^{2+}$ transients across 200 simulations with different random number seeds. This grand-average $Ca^{2+}$ influx time series was used as input for our biochemically detailed single-compartmental model of the post-synaptic spine.

#### Mapping ion-channel encoding genes to modelled current entities

The L23PC model describes the following ionic currents: high-voltage activated $Ca^{2+}$ current ($I_{\mathrm{Ca,HVA}}$), low-voltage activated (LVA) $Ca^{2+}$ current ($I_{\mathrm{Ca,LVA}}$), non-specific cationic current ($I_h$), muscarinic $K^{+}$ current ($I_m$), transient ($I_{\mathrm{K,t}}$) and persistent ($I_{\mathrm{K,p}}$) $K^{+}$ currents, transient ($I_{\mathrm{Na,t}}$) and persistent ($I_{\mathrm{Na,p}}$) $Na^{+}$ currents, $Ca^{2+}$-activated SK current ($I_{\mathrm{SK}}$), Kv3.1-mediated $K^{+}$ current $I_{\mathrm{Kv3.1}}$, and the leak current ($I_{\mathrm{leak}}$). Changes in the expression of the subunits contributing to these currents are modelled by altered maximal conductance ($\bar{g}$) of the underlying current, as typically done in the field (see e.g. [O'Leary et al., 2014, Lemaire et al., 2021]). For many of these currents ($I_{\mathrm{Ca,HVA}}$, $I_{\mathrm{Ca,LVA}}$, $I_h$, $I_m$, and $I_{\mathrm{SK}}$), it is known what the main ion-channel-subunit-encoding genes contributing to these currents are: of the genes listed in Table 1B, CACNA1C and CACNA1D contribute to $I_{\mathrm{Ca,HVA}}$, CACNA1I to $I_{\mathrm{Ca,LVA}}$, HCN1 to $I_h$, and KCNQ3 to $I_m$. However, the description of transient and persistent $Na^+$ currents $I_{\mathrm{Na,t}}$ and $I_{\mathrm{Na,p}}$ (both of which are tetrodotoxin-sensitive [Hamill et al., 1991, Magistretti and Alonso, 1999]) leaves uncertain which subunits or subunit compositions comprise the two $Na^+$ currents. The same can be said of the transient and persistent $K^+$ currents ($I_{\mathrm{K,t}}$ and $I_{\mathrm{K,p}}$) as well as the leak current. We modelled the effects of KCNB1 and KCND3 as altered $I_{\mathrm{K,p}}$ since the experimentally measured kinetics of currents through these genes are closer to the dynamics of $I_{\mathrm{K,p}}$ than those of $I_{\mathrm{K,t}}$ (KCNB1-mediated $K^+$ currents have activation time constant within the range 2–10 ms [Scholle et al., 2004] and KCND3-mediated currents within 1–20 ms [Wang et al., 2004], while the time constants of the modelled currents $I_{\mathrm{K,t}}$ and $I_{\mathrm{K,p}}$ are within the range 0.1–0.4 or 1.4–16.6 ms, respectively). The $\beta$-subunit encoding gene SCN1B contributes positively to Nav1.1-mediated currents [Martinez-Moreno et al., 2020], which have inactivation time constants in the range 1–3 ms [Ghovanloo et al., 2018]. Similarly, the inactivation time constants of SCN9A-mediated currents have been found to lie in the range 0.2–0.5 ms [Zhang et al., 2018] or 0.5–7 ms [Vijayaragavan et al., 2001], although larger (4–20 ms) time constants were also observed if $\beta$ subunits were present [Vijayaragavan et al., 2001]. Here, we consider both SCN1B and SCN9A to affect the transient $Na^+$ current $I_{\mathrm{Na,t}}$ since its dynamics are closer to the reported ranges: $I_{\mathrm{Na,t}}$ has inactivation time constants in the range 0.2–1.9 ms while $I_{\mathrm{Na,p}}$ has the time constants in the range 430–2200 ms. Finally, KCNK4 encodes a subunit (K2p4.1) of a two-pore leak $K^+$ channel, and thus we model the effects of altered KCNK4 expression as changes in $I_{\mathrm{leak}}$.

#### Usability of the model for describing brain area-specific synaptic plasticity

With our previous, less complete model of cortical post-synaptic plasticity we were able to reproduce a large set of experimental results on LTP/LTD and their molecular dependencies by varying the concentration of different proteins available to the spine [Mäki-Marttunen et al., 2020]. However, the model failed to reproduce CaMKII-dependent forms of plasticity in visual cortex as reported in [Kirkwood et al., 1997]. Moreover, the model did not describe the PKC-dependent dynamics of neurogranin, a small protein highly enriched in hippocampus and cortex [Represa et al., 1990], which facilitates LTP by releasing buffered CaM in response to PKC activation [Zhong et al., 2011]. Furthermore, the assumption of spontaneous insertion of non-S845-phosphorylated GluR1 to the membrane hindered the model of [Mäki-Marttunen et al., 2020] from reproducing PKA-dependent LTD observed especially in visual cortex [Fischer et al., 2004, Seol et al., 2007]. Here, we performed multi-objective optimisation where the protein concentrations were varied to fit the LTP/LTD outcome observed in the experiments on cortical synaptic plasticity. Many of the experimental data we used [Ma et al., 2008, Sáez-Briones et al., 2015, Flores et al., 2011, Hardingham et al., 2003, Song et al., 2017, Zhou et al., 2013, Kirkwood et al., 1997, Kotak et al., 2007, Zhong et al., 2011] introduced pharmacological or genetic manipulations of the studied post-synaptic neurons, and thus, similar to [Mäki-Marttunen et al., 2020], we included each manipulation in the optimization. We varied the sizes of glutamatergic and beta-adrenergic ligand fluxes and $Ca^{2+}$ flux and the protein concentrations by $\pm 100\%$. We accepted parameter sets that reproduced the correct LTP/LTD outcome up to an average distance of 1 standard deviation (SD) from that reported in the experimental data, both for the control experiment and pharmacological or genetic manipulations when applicable.

We were able to reproduce all the data (9 data sets) that were successfully reproduced in [Mäki-Marttunen et al., 2020],

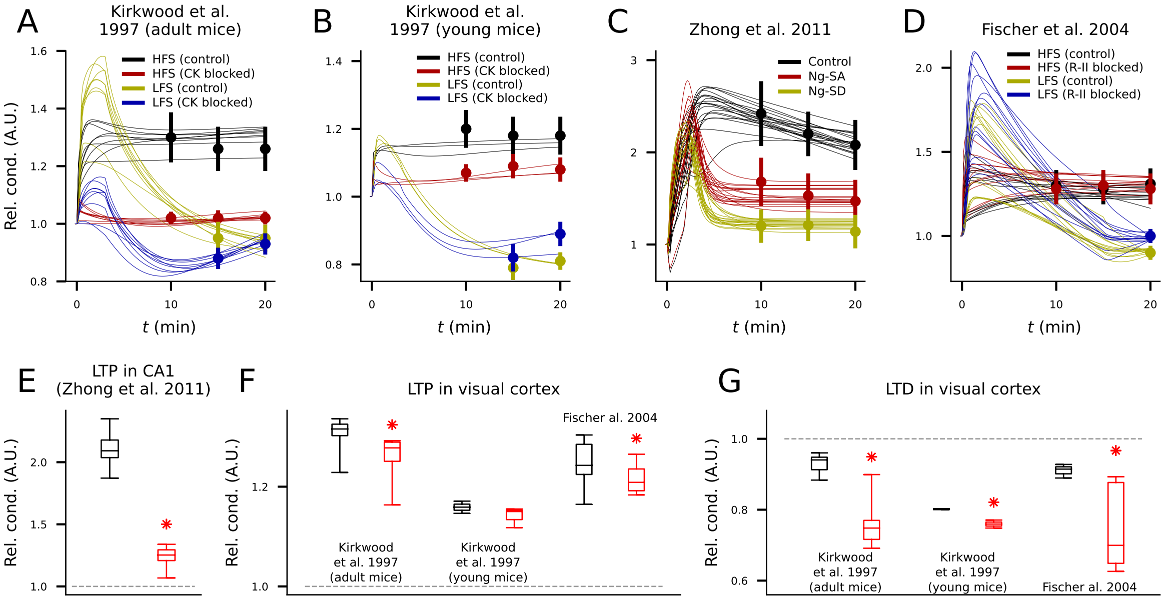


Figure S1: **Reproduction of multi-protocol plasticity experiments and predictions with additional blockers. A–D**: Fit of the model to synaptic plasticity data with pharmacological or genetic manipulations from adult mouse visual cortex [Kirkwood et al., 1997] (A), young mouse visual cortex [Kirkwood et al., 1997] (B), CA1 [Zhong et al., 2011] (C), or mouse V1 [Fischer et al., 2004] (D). Curves represent the predicted time course of the relative synaptic conductance using different acceptable parameter sets — each simulation is repeated with the same parameter set using the 3–4 different conditions (different colors). The vertical bars represent the corresponding mean and SD of each condition from the experimental data. **E–G**: Model predictions with the acceptable parameter sets from the fitting in control experiments (black) and in experiments where all neurogranin is removed (red; E–G). In each panel the simulations were run using the model parameters fitted to the underlying data. The panel (E) shows the model predictions for the protocols of [Zhong et al., 2011], the panel (F) shows those for the LTP-inducing protocols of [Kirkwood et al., 1997] and [Fischer et al., 2004], and the panel (G) shows those for the LTD-inducing protocols of [Kirkwood et al., 1997] and [Fischer et al., 2004]. The bars represent the predicted synaptic conductance (minimum, 25% quartile, median, 75% quartile, and maximum across the parameter sets) 20 min after stimulus onset. Asterisks denote significant differences between the model predictions for blocked and control cases (U-test, $p<0.05$).

and, additionally, CaMKII- and stimulation frequency-dependent plasticity in V1 [Kirkwood et al., 1997] in adult (Fig. S1A) and young (Fig. S1B) animals, effects of PKC-neurogranin interactions on postsynaptic LTP in CA1 [Zhong et al., 2011] (Fig. S1C), and $\beta$-adrenergic receptor and stimulation frequency-dependent plasticity in V1 [Fischer et al., 2004] (Fig. S1D). The parameter values for three best parameter sets for each fitted data set are given in Table S3.

To explore the LTP/LTD mechanisms in the brain regions considered in these experimental studies, we examined the plasticity response under pharmacological blockade. Namely, we used the parameters from the acceptable fits of Fig. S1 and simulated the model under the same stimulation protocol as before but additionally removing neurogranin. Our results suggest that completely removing neurogranin from the synapse (assuming, however, the same amount of CaM) significantly (U-test, $p<0.05$) impairs the CA1 LTP approximately as much as the Ng-SD mutation, which blocks the CaM-binding of neurogranin by making all neurogranin effectively phosphorylated (LTP amplitude +4–34% for complete removal of neurogranin, Fig. S1E; +17–27% for Ng-SD, Fig. S1C). The effects of removal of neurogranin on HFS-induced LTP in visual cortex were milder but qualitatively similar (Fig. S1F), whereas it had an amplifying effect on LFS-induced LTD in visual cortex: our model fit to the data from visual cortex of both adult and young mice [Kirkwood et al., 1997, Fischer et al., 2004] predicted a significantly (U-test, $p<0.05$) enhanced post-20 min LTD compared to controls (Fig. S1G). Taken together, we produced brain area-specific models that can be readily used to explore the role of risk proteins of mental disorders in synaptic plasticity in various cortical areas.

Table S3: Model parameters fit to 13 brain area-specific datasets with acceptable accuracy (1 SD from experimental data). The index column, containing letters A–M, indicates the plasticity data set fitted — these are the following (mostly the same as in [Mäki-Marttunen et al., 2020]): A) Entorhinal cortex, horizontal pathway [Ma et al., 2008], B) Entorhinalcortex, ascending pathway [Ma et al., 2008], C) Prefrontal cortex (CC→PFC synapses) [Sáez-Briones et al., 2015], D) Prefrontal cortex (CC→PFC synapses) [Flores et al., 2011], E) Barrel cortex (L4→L2/3 synapses) [Hardingham et al., 2003], F) Anterior cingulate cortex (L5/6→L2/3 synapses) [Song et al., 2017], G) Prefrontal cortex (L2/3→L2/3 synapses) [Zhou et al., 2013], H) Visual cortex (L4→L3 synapses, mature) [Kirkwood et al., 1997], I) Visual cortex (L4→L3 synapses, young) [Kirkwood et al., 1997], J) Auditory cortex (L6→L5 synapses, LTP-expressing neurons) [Kotak et al., 2007], K) Audirory cortex (L6→L5 synapses, LTD-expressing neurons) [Kotak et al., 2007], L) CA1 (CA3→CA1 synapses) [Zhong et al., 2011], and M) Visual cortex (L4→L2/3 synapses) [Fischer et al., 2004]. Here, three acceptable parameter sets are reported from each data set. The first four columns report the following model parameters 1) $Ca^{2+}$ flux (particles/ms), 2) flux of $\beta$-adrenergic ligand (particles/ms), 3) flux of glutamate to activate mGluRs (particles/ms), and 4) the ratio of GluR1/(GluR1+GluR2). The next two columns are factors for multiple upstream PKC or PKA-pathway proteins, namely, 5) mGluR, M1R, Gq, and PLC, or 6) $\beta$-adrenergic receptor, Gs, AC1 and AC8. The columns 7–19 are factors for single proteins, namely 7) CaM, 8) neurogranin, 9) CaMKII, 10) leak channel, 11) NCX, 12) PKC, 13) PKA, 14) PP1, 15) I1, 16) PP2B, 17) PP2A, 18) PDE1A, and 19) PDE4. In the last data set (M; from [Fischer et al., 2004]), spontaneous S845 phosphorylation of GluR1 receptors was allowed to model non-PKA-mediated S845 phosphorylation. The rate of this spontaneous phosphorylation was fitted along with the other parameters and is given in the 20th column (in 1/(ms)).

| | | | | | | | | | | | | | | | | | | | | |
|---|---|---|---|---|---|---|---|---|---|---|---|---|---|---|---|---|---|---|---|---|
| A | 231.201 | 3.678 | 0.0 | 0.833 | 0.570 | 1.060 | 1.427 | 0.637 | 0.498 | 0.940 | 0.620 | 1.355 | 1.163 | 0.346 | 0.798 | 0.257 | 1.432 | 1.360 | 0.235 | |
| A | 235.738 | 3.652 | 0.633 | 0.831 | 0.553 | 1.194 | 1.361 | 0.617 | 0.501 | 0.974 | 0.592 | 1.335 | 1.114 | 0.330 | 0.904 | 0.258 | 1.424 | 1.409 | 0.286 | |
| A | 235.187 | 3.682 | 0.0 | 0.838 | 0.638 | 1.031 | 1.425 | 0.629 | 0.621 | 1.007 | 0.619 | 1.356 | 1.170 | 0.419 | 0.801 | 0.281 | 1.433 | 1.475 | 0.230 | |
| B | 331.794 | 3.007 | 199.023 | 0.444 | 1.828 | 1.870 | 1.696 | 0.507 | 0.461 | 1.551 | 1.086 | 1.873 | 0.876 | 1.607 | 0.918 | 1.293 | 1.244 | 0.085 | 0.813 | |
| B | 322.542 | 4.199 | 177.723 | 0.331 | 1.812 | 1.542 | 1.949 | 0.581 | 0.444 | 1.645 | 1.313 | 1.864 | 0.349 | 1.469 | 1.448 | 0.916 | 1.010 | 0.278 | 1.228 | |
| B | 365.428 | 3.902 | 177.292 | 0.355 | 1.698 | 1.543 | 1.830 | 0.512 | 1.022 | 1.618 | 1.293 | 1.957 | 0.389 | 1.563 | 1.225 | 1.065 | 1.211 | 0.376 | 1.246 | |
| C | 132.842 | 3.456 | 24.339 | 0.313 | 0.392 | 1.373 | 1.879 | 1.335 | 0.678 | 1.971 | 0.551 | 0.516 | 0.169 | 1.350 | 1.539 | 0.124 | 0.073 | 0.844 | 0.428 | |
| C | 166.100 | 3.457 | 178.802 | 0.794 | 1.431 | 1.604 | 2.000 | 1.229 | 0.666 | 1.153 | 1.934 | 1.106 | 0.455 | 0.433 | 0.905 | 0.563 | 0.953 | 1.259 | 1.323 | |
| C | 108.171 | 1.436 | 100.173 | 0.835 | 0.315 | 1.389 | 1.872 | 1.417 | 0.018 | 1.987 | 1.051 | 0.830 | 0.334 | 0.372 | 1.518 | 1.901 | 1.410 | 1.040 | 1.346 | |
| D | 146.253 | 2.598 | 60.755 | 0.660 | 2.000 | 2.000 | 0.901 | 0.877 | 1.175 | 0.891 | 0.525 | 1.405 | 0.446 | 1.939 | 1.446 | 0.249 | 1.110 | 0.885 | 1.941 | |
| D | 135.773 | 2.574 | 60.781 | 0.643 | 1.908 | 2.000 | 0.881 | 0.882 | 1.184 | 0.879 | 0.289 | 1.327 | 0.594 | 1.897 | 1.473 | 0.214 | 0.903 | 0.975 | 1.873 | |
| D | 151.399 | 3.274 | 96.919 | 0.800 | 1.513 | 1.417 | 0.956 | 0.992 | 0.891 | 0.702 | 0.462 | 0.205 | 0.421 | 0.342 | 0.549 | 1.053 | 1.132 | 0.777 | 0.768 | |
| E | 509.696 | 5.000 | 89.190 | 0.491 | 0.606 | 1.495 | 1.568 | 1.155 | 1.664 | 1.574 | 0.00749 | 0.0 | 1.100 | 1.731 | 1.366 | 1.912 | 1.141 | 0.508 | 0.587 | |
| E | 293.219 | 4.827 | 65.572 | 0.398 | 0.670 | 1.433 | 1.604 | 1.184 | 1.670 | 1.495 | 0.042 | 0.104 | 1.227 | 1.542 | 1.338 | 1.957 | 1.113 | 0.445 | 0.314 | |
| E | 641.407 | 4.642 | 58.121 | 0.412 | 0.764 | 1.503 | 1.646 | 1.024 | 1.422 | 1.505 | 0.0 | 0.000664 | 1.231 | 1.676 | 1.313 | 2.000 | 1.078 | 0.584 | 0.233 | |
| F | 129.309 | 4.573 | 91.454 | 0.949 | 0.935 | 1.618 | 1.038 | 0.786 | 0.245 | 1.900 | 0.909 | 1.092 | 1.273 | 1.448 | 0.297 | 1.488 | 0.0 | 0.581 | 0.629 | |
| F | 128.115 | 5.000 | 116.861 | 0.904 | 0.968 | 1.439 | 0.992 | 0.789 | 0.224 | 1.931 | 0.927 | 1.070 | 1.088 | 1.266 | 0.323 | 1.412 | 0.0 | 0.503 | 0.599 | |
| F | 130.783 | 4.389 | 89.165 | 0.939 | 0.940 | 1.581 | 1.088 | 0.786 | 0.241 | 1.927 | 0.939 | 1.227 | 1.475 | 1.428 | 0.378 | 1.484 | 0.0 | 0.584 | 0.617 | |
| G | 579.195 | 3.570 | 118.880 | 0.644 | 1.444 | 0.311 | 1.960 | 1.950 | 0.377 | 2.000 | 1.030 | 1.046 | 1.776 | 1.707 | 2.000 | 1.731 | 1.285 | 0.157 | 1.090 | |
| G | 251.668 | 3.557 | 119.101 | 0.527 | 1.445 | 0.304 | 2.000 | 1.964 | 0.373 | 2.000 | 1.122 | 1.011 | 1.516 | 1.546 | 1.997 | 1.834 | 1.278 | 0.133 | 1.177 | |
| G | 303.924 | 3.893 | 162.119 | 0.714 | 1.381 | 0.266 | 1.819 | 1.795 | 0.186 | 1.970 | 1.020 | 1.160 | 1.748 | 1.561 | 2.000 | 1.721 | 1.469 | 0.131 | 1.147 | |
| H | 576.135 | 4.215 | 103.839 | 0.685 | 1.344 | 1.662 | 1.935 | 1.831 | 1.153 | 1.076 | 1.039 | 0.217 | 1.407 | 0.207 | 1.006 | 0.335 | 1.574 | 1.166 | 0.362 | |
| H | 603.773 | 1.439 | 61.090 | 0.714 | 1.949 | 1.856 | 1.492 | 1.747 | 1.955 | 1.267 | 1.531 | 0.040 | 1.290 | 1.054 | 1.207 | 0.146 | 1.035 | 1.015 | 1.390 | |
| H | 575.618 | 4.138 | 101.016 | 0.683 | 1.330 | 1.655 | 1.935 | 1.832 | 1.308 | 1.064 | 1.164 | 0.264 | 1.428 | 0.234 | 1.019 | 0.343 | 1.570 | 1.163 | 0.416 | |
| I | 597.481 | 0.349 | 70.615 | 0.858 | 1.961 | 1.874 | 1.976 | 0.722 | 2.000 | 1.321 | 1.307 | 0.199 | 1.606 | 0.0 | 1.081 | 0.695 | 0.132 | 1.246 | 1.355 | |
| I | 688.112 | 1.948 | 29.187 | 0.908 | 1.652 | 1.069 | 1.547 | 1.632 | 0.446 | 1.912 | 1.531 | 1.066 | 1.780 | 0.023 | 0.948 | 0.114 | 1.675 | 0.939 | 0.481 | |
| I | 661.438 | 0.160 | 118.410 | 0.808 | 1.627 | 1.223 | 1.823 | 1.117 | 0.630 | 2.000 | 2.000 | 1.837 | 1.406 | 0.040 | 1.780 | 0.958 | 1.467 | 0.980 | 1.305 | |
| J | 799.265 | 2.633 | 33.622 | 0.588 | 1.959 | 1.871 | 0.672 | 0.678 | 0.984 | 0.871 | 1.355 | 1.531 | 0.214 | 0.406 | 0.907 | 1.500 | 0.446 | 0.127 | 1.427 | |
| J | 604.315 | 2.275 | 116.522 | 0.607 | 0.122 | 1.518 | 1.253 | 1.405 | 0.810 | 0.802 | 0.633 | 0.454 | 1.859 | 0.940 | 0.595 | 0.396 | 0.174 | 0.326 | 1.486 | |
| J | 583.887 | 1.615 | 114.658 | 0.615 | 0.141 | 1.337 | 1.262 | 1.485 | 0.871 | 0.778 | 0.532 | 0.422 | 1.757 | 0.616 | 0.555 | 0.401 | 0.211 | 0.327 | 1.498 | |
| K | 457.679 | 1.141 | 196.536 | 0.916 | 0.129 | 1.599 | 1.116 | 0.898 | 1.218 | 1.026 | 1.372 | 1.494 | 1.046 | 1.624 | 1.935 | 1.183 | 0.729 | 1.112 | 0.557 | |
| K | 395.217 | 0.0 | 194.514 | 0.915 | 0.104 | 1.579 | 1.080 | 0.879 | 1.391 | 1.002 | 1.020 | 1.109 | 0.833 | 1.486 | 1.897 | 1.082 | 0.890 | 1.072 | 0.405 | |
| K | 454.349 | 0.355 | 184.717 | 0.938 | 0.00187 | 1.660 | 1.085 | 0.859 | 1.192 | 0.945 | 1.266 | 1.124 | 0.990 | 1.669 | 1.881 | 1.016 | 0.712 | 1.130 | 0.550 | |
| L | 521.028 | 4.477 | 169.096 | 0.291 | 0.229 | 1.984 | 0.384 | 0.492 | 1.257 | 0.580 | 0.031 | 1.756 | 1.310 | 1.532 | 0.918 | 1.270 | 2.000 | 0.983 | 1.143 | |
| L | 462.059 | 4.099 | 187.155 | 0.395 | 0.210 | 2.000 | 0.559 | 1.021 | 1.271 | 0.533 | 0.019 | 1.211 | 1.382 | 1.389 | 0.895 | 1.523 | 2.000 | 0.822 | 0.979 | |
| L | 423.226 | 4.140 | 176.909 | 0.337 | 0.221 | 1.882 | 0.396 | 0.693 | 1.335 | 0.562 | 0.0 | 1.266 | 1.248 | 1.688 | 0.922 | 1.436 | 1.990 | 0.865 | 0.823 | |
| M | 1378.800 | 2.798 | 43.794 | 0.655 | 0.382 | 0.291 | 1.927 | 1.343 | 0.811 | 0.941 | 0.157 | 0.223 | 0.456 | 0.169 | 1.672 | 1.169 | 0.396 | 0.424 | 1.167 | $2.435 \cdot 10^{-7}$ |
| M | 1459.975 | 2.736 | 62.952 | 0.620 | 0.480 | 0.106 | 1.981 | 1.018 | 0.801 | 0.793 | 0.368 | 0.086 | 0.481 | 0.047 | 1.543 | 0.674 | 0.458 | 0.337 | 1.456 | $4.888 \cdot 10^{-7}$ |
| M | 1732.295 | 2.568 | 53.922 | 0.677 | 0.240 | 0.286 | 2.000 | 1.260 | 0.818 | 0.817 | 0.304 | 0.383 | 0.410 | 0.121 | 1.725 | 1.252 | 0.499 | 0.450 | 1.493 | $4.573 \cdot 10^{-7}$ |

Table S4: List of genes considered in the genetic analyses.

- Genes regulating plasticity (not exclusively through PKA or PKC pathway)
- Genes regulating plasticity through PKA pathway
- Genes regulating plasticity through PKC pathway
- Genes encoding ion-channels and calcium transporters, excluding genes belonging to plasticity-regulating gene sets

CACNA1A
CACNA1B
CACNA1C
CACNA1D
CACNA1E
CACNA1F
CACNA1G
CACNA1H
CACNA1I
CACNA1S
CACNB1
CACNB2
CACNB3
CACNB4
CACNA2D1
CACNA2D2
CACNA2D3
CACNA2D4
CACNG1
CACNG2
CACNG3
CACNG4
CACNG5
CACNG6
CACNG7
CACNG8
SCN1A
SCN2A
SCN3A
SCN4A
SCN5A
SCN7A
SCN8A
SCN9A
SCN10A
SCN11A
SCN1B
SCN2B
SCN3B
SCN4B
KCNN1
KCNN2
KCNN3
KCNN4
KCNA1
KCNA2
KCNA3
KCNA4
KCNA5
KCNA6
KCNA7
KCNA10
KCNB1
KCNB2
KCNC1
KCNC2
KCNC3
KCNC4
KCND1
KCND2
KCND3
KCNQ1
KCNQ2
KCNQ3
KCNQ4
KCNQ5
KCNH1
KCNH2
KCNH3
KCNH4
KCNH5
KCNH6
KCNH7
KCNH8
KCNF1
KCNG1
KCNG2
KCNG3
KCNG4
KCNJ1
KCNJ2
KCNJ3
KCNJ4
KCNJ5
KCNJ6
KCNJ8
KCNJ9
KCNJ10
KCNJ11
KCNJ12
KCNJ13
KCNJ14
KCNJ15
KCNJ16
KCNJ17
KCNK1
KCNK2
KCNK3
KCNK4
KCNK5
KCNK6
KCNK7
KCNK9
KCNK10
KCNK12
KCNK13
KCNK15
KCNK16
KCNK17
KCNK18
KCNS1
KCNS2
KCNS3
KCNT1
KCNT2

KCNU1
KCNMA1
KCNMB1
KCNMB2
KCNMB3
KCNMB4
KCNV1
KCNV2
KCNAB1
KCNAB2
KCNAB3
KCNE1
KCNE2
KCNE3
KCNE4
KCNE1L
KCNIP1
KCNIP2
KCNIP3
KCNIP4
HCN1
HCN2
HCN3
HCN4
NKAIN1
NKAIN2
NKAIN3
NKAIN4
NKAIN1P1
NKAIN1P2
ATP2A1
ATP2A2
ATP2A3
ATP2B1
ATP2B2
ATP2B3
ATP2B4
ATP2C1
ATP2C2
ITPR1
ITPR2
ITPR3
ITPRIP
SLC8A1
SLC8A2
SLC8A3
SLC8B1
SLC24A1
SLC24A2
SLC24A3
SLC24A4
SLC24A5
SLC32A1
SLC44A1
SLC44A2
SLC44A3
SLC44A4
SLC44A5
SLC44A3-AS1
SLC44A3P1
SLC12A1
SLC12A2
SLC12A3
SLC12A4
SLC12A5
SLC12A6
SLC12A7
SLC12A8
SLC12A9
SLC12A5-AS1
SLC12A9-AS1
CATSPER1
CATSPER2
CATSPER2P1
CATSPER3
CATSPER4
CATSPERB
CATSPERD
CATSPERE
CATSPERG
CATSPERZ
TRPC1
TRPC2
TRPC3
TRPC4
TRPC5
TRPC6
TRPC7
TRPC8
TRPC4AP
TRPC4OS
TRPC7-AS1
TRPC7-AS2
TRPM1
TRPM2
TRPM3
TRPM4
TRPM5
TRPM6
TRPM7
TRPM8
TRPV1
TRPV2
TRPV3
TRPV4
TRPV5
TRPV6
GRIA1
GRIA2
GRIA3
GRIA4
GRID1
GRID2
GRID1-AS1
GRID2IP

GRIN1
GRIN2A
GRIN2B
GRIN2C
GRIN2D
GRIN3A
GRIN3B
GCOM2
GABRA1
GABRA2
GABRA3
GABRA4
GABRA5
GABRA6
GABRB1
GABRB2
GABRB3
GABRD
GABRE
GABRG1
GABRG2
GABRG3
GABRG3-AS1
GABRP
GABRQ
GABRR1
GABRR2
GABRR3
GABBR1
GABBR2
CHRNA1
CHRNA2
CHRNA3
CHRNA4
CHRNA5
CHRNA6
CHRNA7
CHRNA9
CHRNA10
CHRNB1
CHRNB2
CHRNB3
CHRNB4
CHRND
CHRNE
CHRNG
GRM1
GRM2
GRM3
GRM4
GRM5
PRKACA
PRKACB
PRKACG
PRKAR1A
PRKAR1B
PRKAR2A
PRKAR2B
AKAP1
AKAP2
AKAP3
AKAP4
AKAP5
AKAP6
AKAP7
AKAP8
AKAP9
AKAP10
AKAP11
AKAP12
AKAP13
AKAP14
AKAP15
AKAP17A
AKAP17ABP
SPHKAP
PRKCA
PRKCB
PRKCD
PRKCE
PRKCG
PRKCH
PRKCI
PRKCQ
PRKCZ
PRKD1
PRKD2
PRKD3
PRKG1
PRKG2
CAMK2A
CAMK2B
CAMK2D
CAMK2G
ADCY1
ADCY2
ADCY3
ADCY4
ADCY5
ADCY6
ADCY7
ADCY8
ADCY9
ADCY10
PLCB1
PLCB2
PLCB3
PLCB4
PLCD1
PLCD3
PLCD4
PLCE1
PLCG1
PLCG2
PLCH1

PLCH2
PLCL1
PLCL2
CALM1
CALM2
CALM3
CALN1
GNAS
GNAS-AS1
GNAL
GNAQ
GNA11
GNA14
GNA14-AS1
GNA15
GNA12
GNA13
GNAI1
GNAI2
GNAI3
GNAO1
GNAZ
GNB1
GNB2
GNB3
GNB4
GNB5
GNB1L
GNG2
GNG3
GNG4
GNG5
GNG7
GNG8
GNG10
GNG11
GNG12
GNG13
GNG14
RGS1
RGS2
RGS3
RGS4
RGS5
RGS6
RGS7
RGS7BP
RGS8
RGS9
RGS9BP
RGS10
RGS11
RGS12
RGS13
RGS14
RGS16
RGS17
RGS18
RGS19
RGS20
RGS21
RGS22
GPSM1
GPSM2
GPSM3
GPS1
GPS2
GPS2P1
GPS2P2
CREB1
ATF4
CREB3
CREB5
CREB3L1
CREB3L2
CREB3L3
CREB3L4
CREBL2
ANP32A
ANP32B
ANP32C
ANP32D
ANP32E
ANP32A-IT1
ANP32AP1
ANP32BP2
ANP32BP3
ADRB1
ADRB2
ADRB3
ADRA1A
ADRA1B
ADRA1D
ADRA2A
ADRA2B
ADRA2C
DRD1
DRD2
DRD3
DRD4
DRD5
HTR1A
HTR1B
HTR1D
HTR1DP1
HTR1E
HTR1F
HTR2A
HTR2B
HTR2C
HTR3A
HTR3B
HTR3C
HTR3D
HTR3E

HTR4
HTR5A
HTR6
HTR7
OPRD1
OPRK1
OPRL1
OPRM1
OPRPN
CHRM1
CHRM2
CHRM3
CHRM4
CHRM5
CALB1
CALB2
S100G
RAPGEF3
NRGN
PPP1R1A
PPP1R1B
PPP1R1C
PPP1CA
PPP1CB
PPP1CC
PPP1R2A
PPP1R2B
PPP1R2C
PPP1R3A
PPP1R3B
PPP1R3C
PPP1R3D
PPP1R3E
PPP1R3F
PPP1R3G
PPP1R7
PPP1R8
PPP1R9A
PPP1R9B
PPP1R10
PPP1R11
PPP1R12A
PPP1R12B
PPP1R12C
PPP1R13B
PPP1R13L
PPP1R14A
PPP1R14B
PPP1R14C
PPP1R14D
PPP1R15A
PPP1R15B
PPP1R16A
PPP1R16B
PHACTR1
PHACTR2
PHACTR3
PHACTR4
PDE1A
PDE1B
PDE1C
PDE2A
PDE3A
PDE3B
PDE4A
PDE4B
PDE4B-AS1
PDE4C
PDE4D
PDE4DIP
PDE4DIPP1
PDE5A
PDE6A
PDE6B
PDE6C
PDE6D
PDE6G
PDE6H
PDE7A
PDE7B
PDE8A
PDE8B
PDE9A
PDE10A
PDE11A
PPP2CA
PPP2CB
PPP2R1A
PPP2R1B
PPP2R2A
PPP2R2B
PPP2R2C
PPP2R2D
PPP2R3A
PPP2R3B
PPP2R3C
PPP2R4
PPP2R5A
PPP2R5B
PPP2R5C
PPP2R5D
PPP2R5E
PPP3CA
PPP3CB
PPP3CC
PPP3R1
PPP3R2
DGKA
DGKB
DGKD
DGKE
DGKG
DGKH
DGKI
DGKK

DGKQ
DGKZ
PLA2R1
PLA2G2A
PLA2G2C
PLA2G2D
PLA2G2E
PLA2G2F
PLA2G3
PLA2G4A
PLA2G4B
PLA2G4C
PLA2G4D
PLA2G4E
PLA2G4F
PLA2G5
PLA2G6
PLA2G7
PLA2G10
PLA2G12A
PLA2G12B
PLA2G15
PITPNM1
PITPNM2
PITPNM3
PITPNA
PITPNB
PITPNC1
NBEA
NBEAL1
NBEAL2
NBEAP1
NBEAP2
NBEAP3
NBEAP4
NBEAP5
NBEAP6
CSNK1A1
CSNK1A1L
CSNK1A1P1
CSNK1A1P2
CSNK1A1P3
CSNK1D
CSNK1E
CSNK1G1
CSNK1G2
CSNK1G3
CSNK1G2-AS1
CSNK1G2P1
CSNK2A1
CSNK2A2
CSNK2A3
CSNK2B
CSNKAIP
CDK5
CDK5R1
CDK5R2
CDK5RAP1
CDK5RAP2
CDK5RAP3
CDK5P1
CLK1
CLK2
CLK3
CLK4
CLK2P1
CLK3P1
CLK3P2
SCGN
NUCB1
NUCB2
NUCB1-AS1
AKT1
AKT2
AKT3
AKTIP
AKT1S1
CLTA
CLTB
CLTC
CLTCL1
CLTRN
SYNGAP1
DLG1
DLG2
DLG3
DLG4
DLG5
MPP
DLGAP5
DLGAP1
DLGAP2
DLGAP3
DLGAP4
DLG1-AS1
DLG2-AS2
DLG3-AS1
DLG5-AS1
SRR
OPCML
OPCML-IT1
OPCML-IT2
NOS1
NOS2
NOS3
NOSIP
NOSTRIN
NOS1AP
NOS2P1
NOS2P2
NOS2P3
NOS2P4

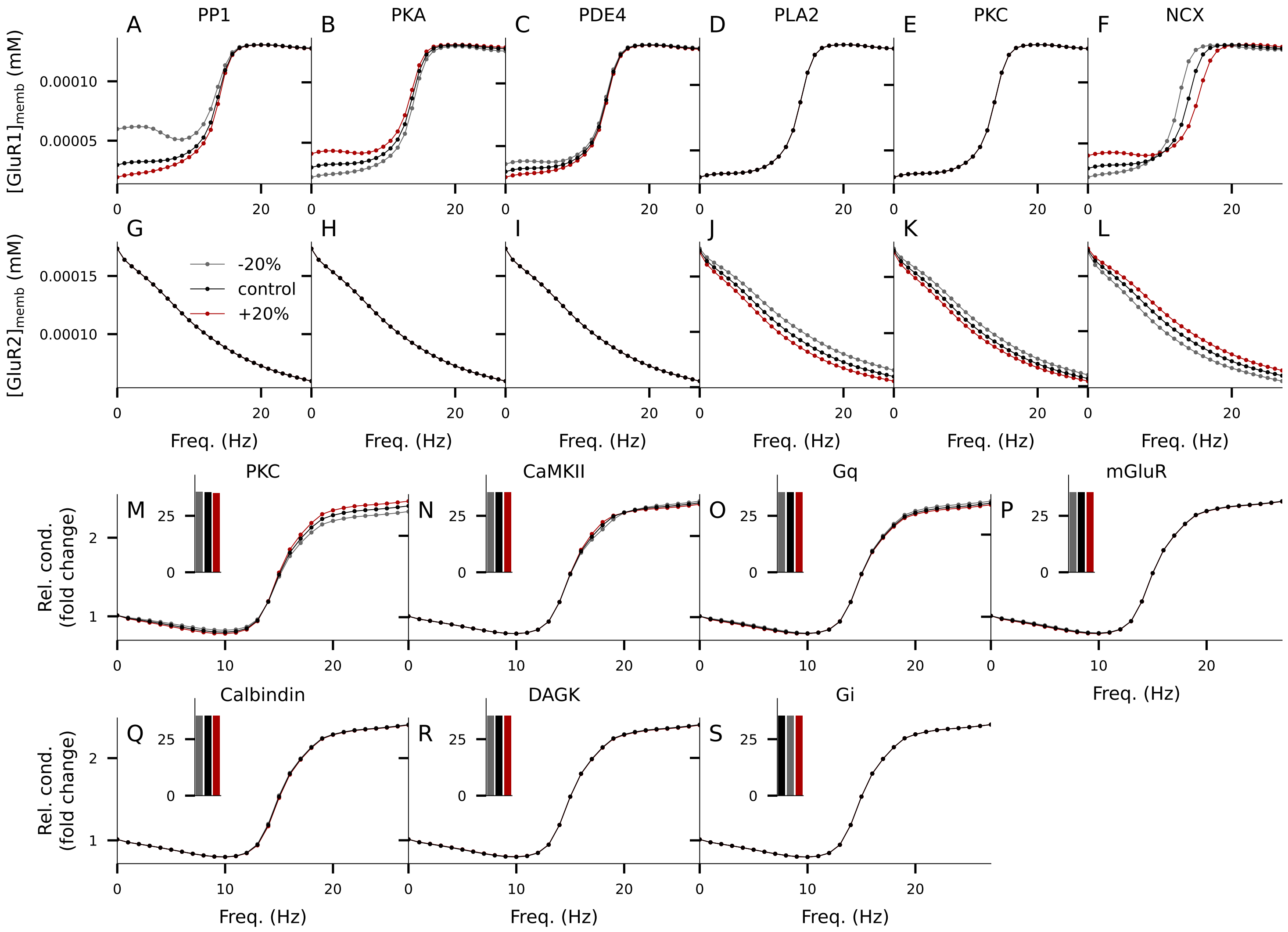


Figure S2: **Complementary analyses of the effects of risk proteins on LTP/LTD curves and how they are mediated by membrane insertion of GluR1 and GluR2. A–L**: The concentration of membrane-inserted GluR1 (A–F) or GluR2 (G–L) receptors on the spine membrane 30 min after stimulus onset as a function of input frequency, given a fixed stimulation time of T=100 s. The red and gray curves represent the data from simulations where the protein concentration of PP1 (A,G), PKA (B,H), PDE4 (C,I), PLA2 (D,J), PKC (E,K), or NCX (F,L) was 20% higher (red) or lower (gray) than in the default model (black). **M–S**: The experiment of Fig. 2A–F repeated for the remaining risk proteins PKC (M), CaMKII (N), Gq (O), mGluR (P), calbindin (Q), DAGK (R), and Gi (S). The effect of ±20% expression of PKC was similar to those of PLA2 (Fig. 2D, while the effects of ±20% expression of CaMKII and Gq were small and those of mGluR, calbindin, DAGK, and Gi were negligible.

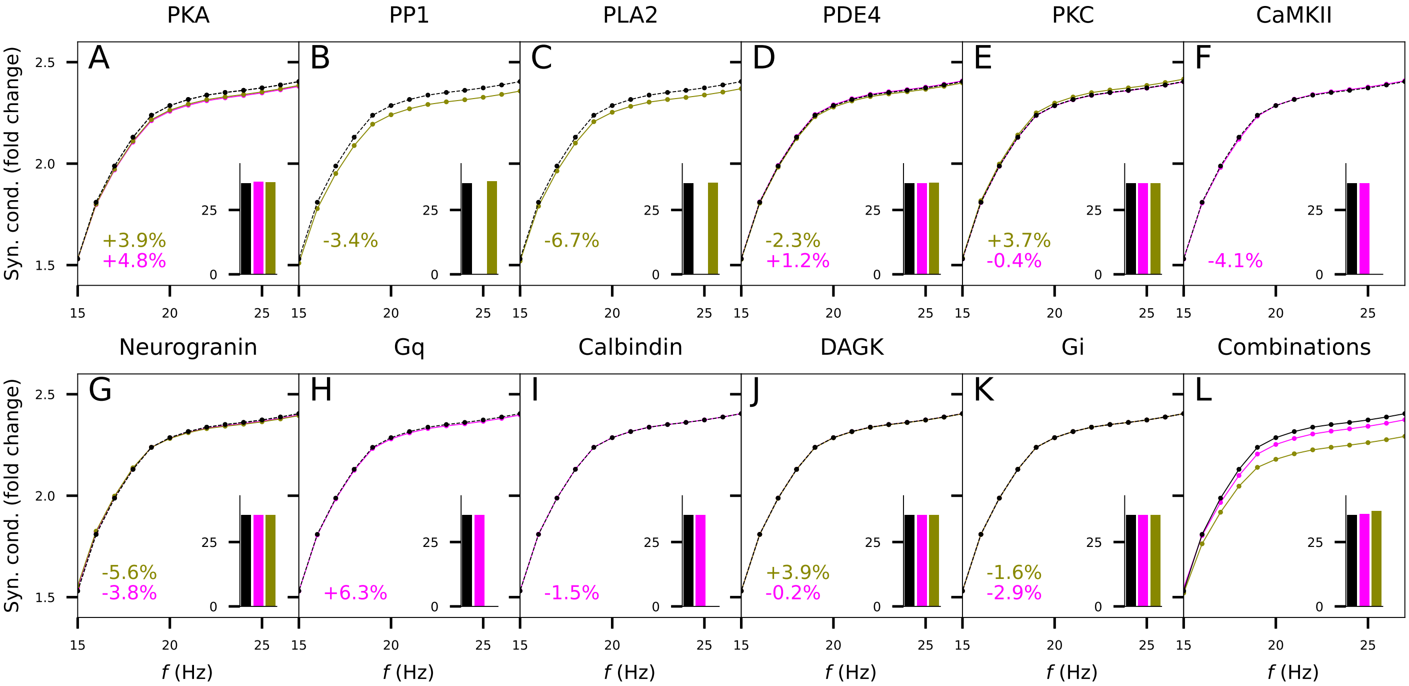


Figure S3: **Effects of SCZ-associated alterations in expression of single proteins, as measured in PFC and ACC, on plasticity. A–K**: The experiments of Fig. 2 were repeated by using the amplitudes of the parameter change obtained from the CommonMind data from PFC (magenta) or ACC (green) instead of the ±20% alteration used in Fig. 2A–F and S2M–S. The implemented changes of protein concentration are printed in the lower-left corner of the panels. The alterations of mGluR and NCX were not included — for the risk genes encoding these proteins the neuron imputation was not successful and thus no data were available to determine the parameter changes. **L**: Zoom-in on high-frequency part of Fig. 2G. The insets show the baseline conductances (pS).

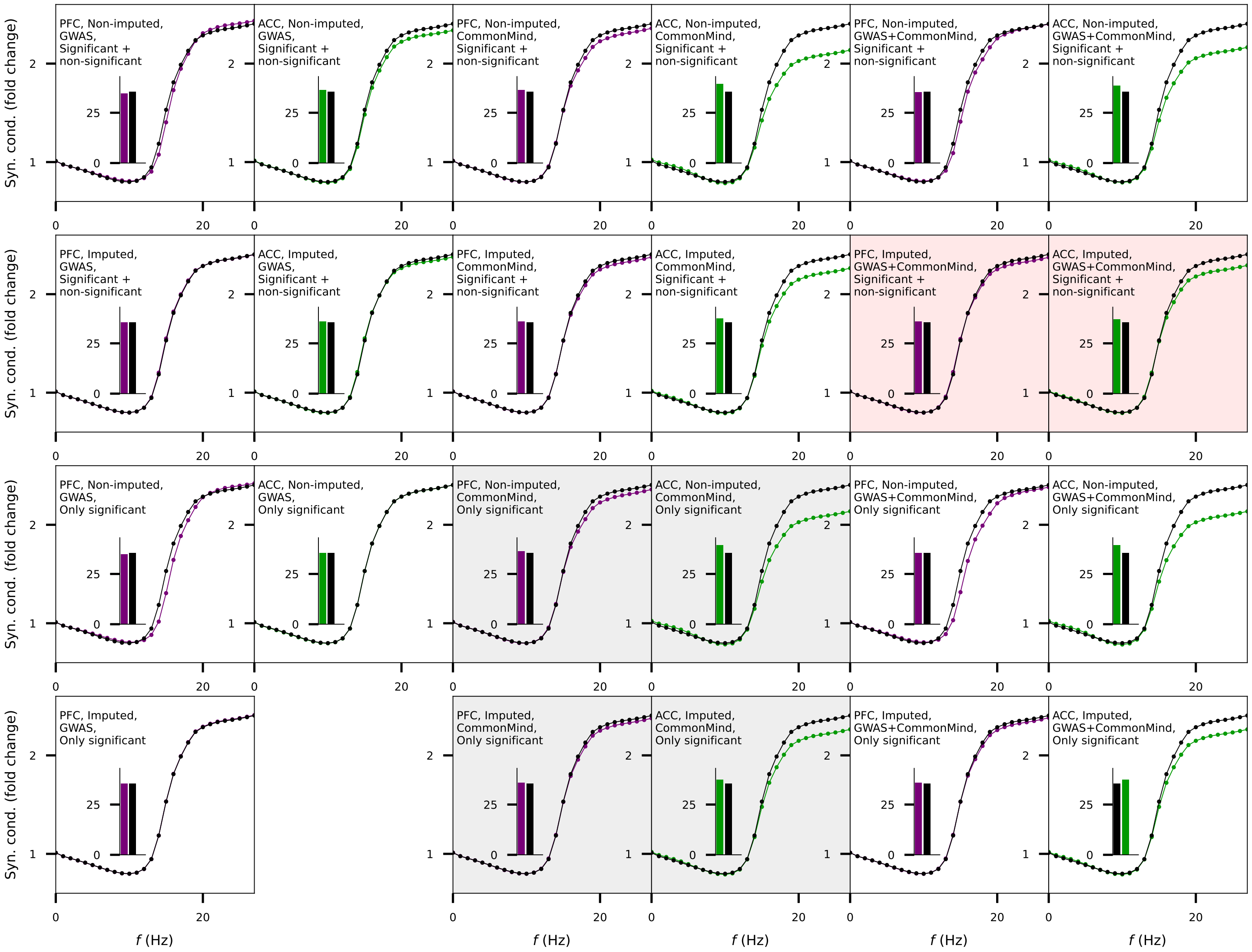


Figure S4: **Effects of combinations of SCZ-associated alterations, as measured in PFC and ACC, on plasticity.** The experiments of Fig. 2G were repeated by using different strategies for determining the parameter changes. First, we used either non-imputed (1st and 3rd rows) or imputed (2nd and 4th rows) data. Second, we identified the genes based either on their minimal p-value in the SCZ GWAS data ($p < 5 \cdot 10^{-6}$; 1st and 2nd columns) or on a strong statistical significance of the difference in expression between SCZ and HC populations in the CommonMind data ($p<0.0005$; 3rd and 4th columns), or we used the union of the two sets of genes (5th and 6th columns). Third, we either used all these genes (1st and 2nd rows) or only those that showed at least moderate ($p<0.05$) statistical significance of difference in expression between SCZ and HC populations in the CommonMind data (3rd and 4th rows). Combinations of these three choices yielded 12 different strategies. In one of these strategies (imputed data, identification based on GWAS, only genes with significantly different expression), there were no genes that were both succesfully imputed and showed significantly different expression between SCZ and HC population, and thus this panel is left blank. The four panels with gray background are redundant: these strategies resulted in the exact same genes and same parameter alterations as the four strategies above since the genes were selected only based on the significance of differential expression in the CommonMind data. The two panels with dim red background correspond to the data of Fig. 2G. The impairment of LTP as observed in Fig. 2G was very robust: apart from two strategies where genes were selected based on GWAS data only and statistical signifance of the differential expression, where there were either no genes to be tested (imputed data) or the LTP was unaltered (non-imputed data), all remaining eight strategies yielded a prediction of decreased LTP amplitude. The predictions for PFC-like alterations were also fairly robust: six out of ten strategies suggested impaired LTP, two suggested an unaltered LTP amplitude, and two suggested decreased LTP amplitude for medium-high stimulus frequencies (approximately 10–20 Hz) and increased LTP amplitude for high frequencies (>20 Hz).

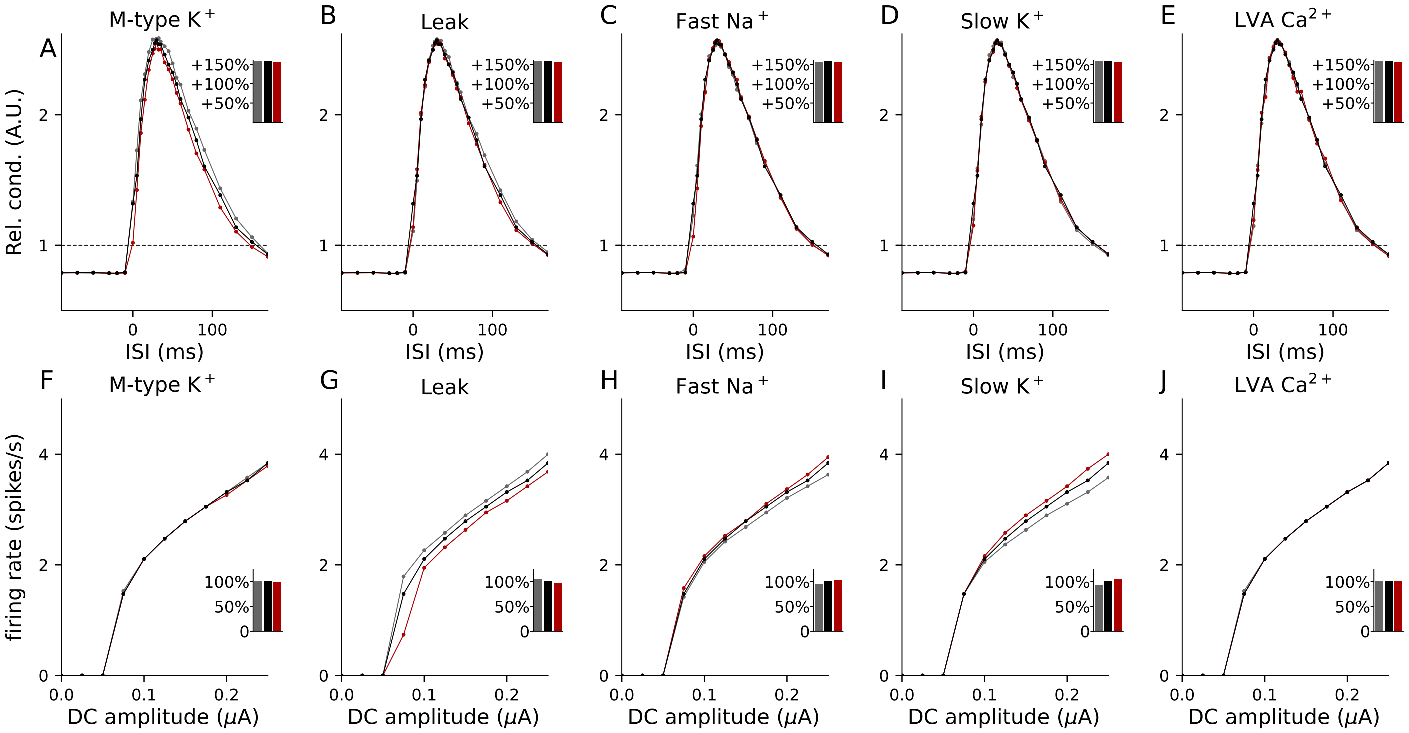


Figure S5: **Complementary analyses of the effects of risk proteins on STDP.** The experiments of Fig. 3D–E and G–H repeated for remaining ion channel-encoding risk genes of SCZ, namely, KCNQ3 (M-type $K^+$ channel, panels A and F), KCNK4 (leak channel, panels B and G), SCN1B and SCN9A (fast $Na^+$ channel, panels C and H), KCNB1 and KCND3 (slow $K^+$ channel, panels D and I), and CACNA1I (LVA $Ca^{2+}$ channel, panels E and J).

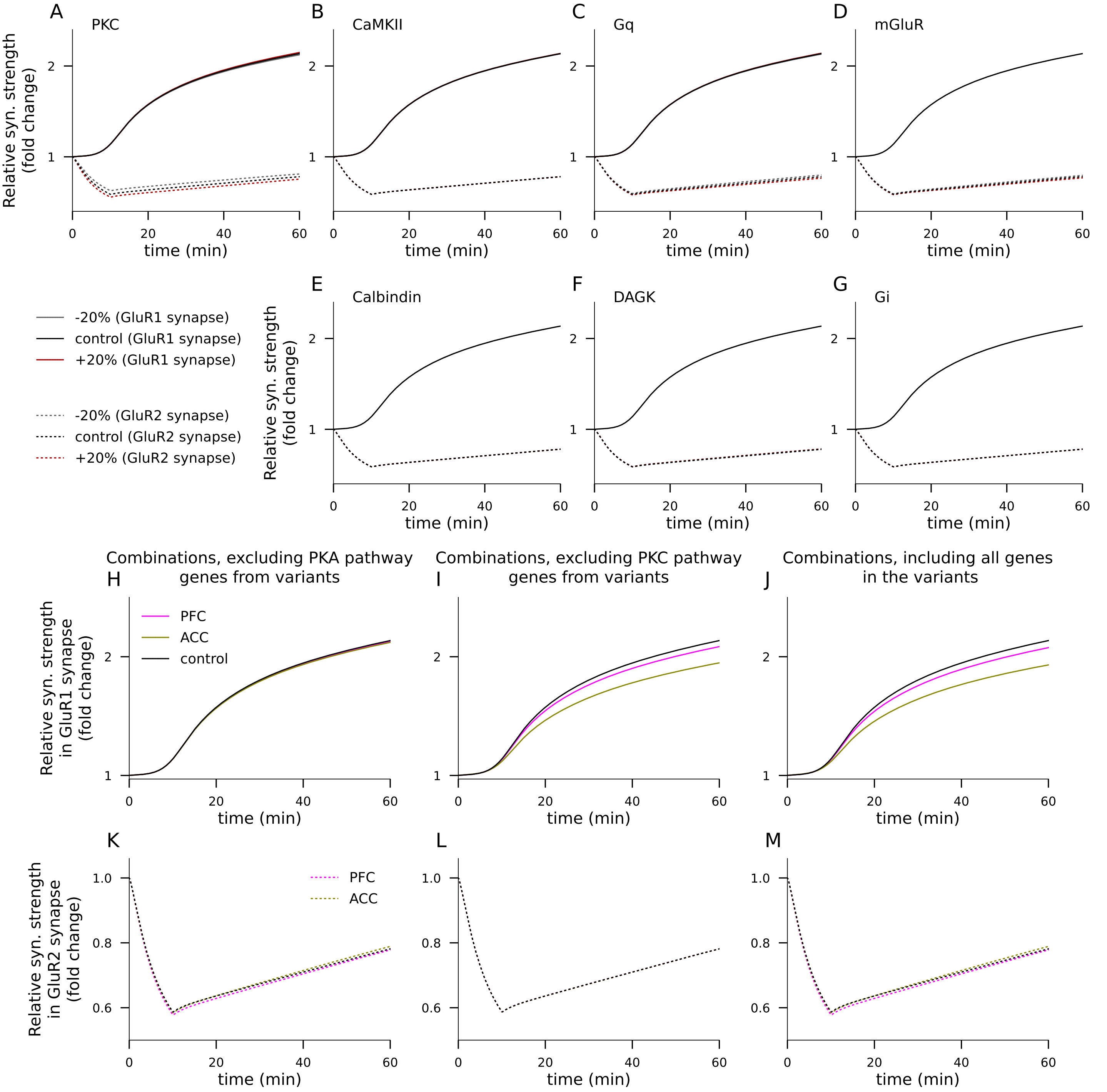


Figure S6: **Complementary analyses of the effects of risk proteins on VEP modulation and the effects of PKA- or PKC-pathway-restricted variants. A–G**: The experiments of Fig. 4E–J repeated for remaining synaptic risk proteins of SCZ, namely, PKC (A), CaMKII (B), Gq (C), mGluR (D), calbindin (E), DAGK (F), and Gi (G). The effects of ±20% expression of PKC were similar to those of PLA2 (Fig. 4H), while the effects of ±20% expression of Gq and mGluR were small and those of CaMKII, calbindin, DAGK, and Gi were negligible. **H–M**: Exclusion of PKA-pathway genes from the combination variants abolished the LTP-impairing effects of the variants in GluR1 synapses and exclusion of PKC-pathway genes abolished the LTD-altering effects of the variants in GluR2 synapses in response to the VEP-like stimulus protocol. By contrast, the combinations where PKA-pathway or PKC-pathway genes were excluded produced similar plasticity in to the VEP-like stimulus protocol in GluR2- or GluR1-type synapses, respectively, as the combinations where all genes were included. Panels (H)–(J) show the relative synaptic conductance in response to the VEP-like stimulus protocol in GluR1-type synapse, whereas panels (K)–(M) show the relative synaptic conductance in response to the VEP-like stimulus protocol in GluR2-type synapse. In (H) and (K), the PKA-pathway gene variants (namely, alterations of parameters [PP1], [PDE4], [Gi], and [PKA]) are left out from the variant combination, whereas in (I) and (L), the PKC-pathway gene variants (namely, alterations of parameters [DAGK], [Ng], [mGluR], [PKC], [Gabgq], and [PLA]) are left out from the variant combination. Panels (J) and (M) show the model predictions for variants where all parameter alterations are included for reference (data of Fig. 4K replotted).

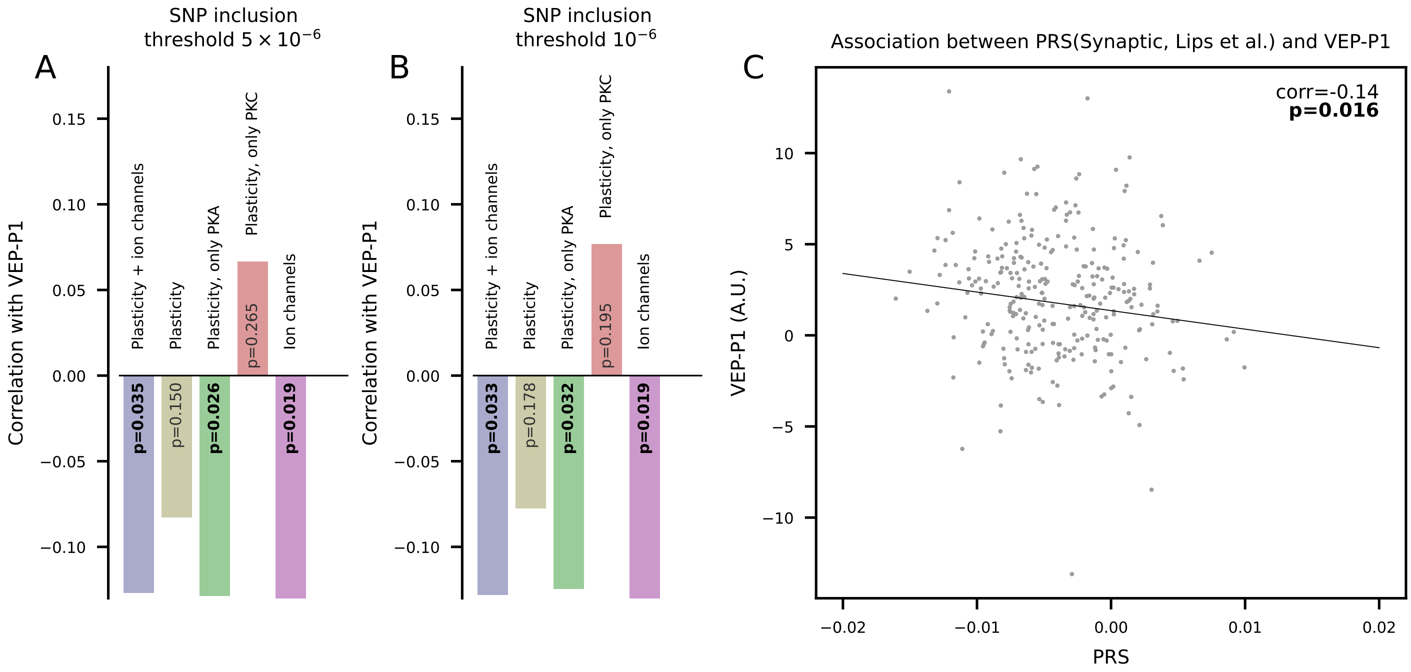


Figure S7: **The genotype-phenotype findings are robust to changes in SNP threshold and the choice of synaptic gene set. A–B**: The experiment of Fig. 5B was repeated by using alternative PRSs where the SNPs were included based on stricter p-value threshold $5 \cdot 10^{-6}$ (A) or $10^{-6}$ (B). **C**: The experiment of Fig. 5D was repeated using an alternative set of synaptic genes, namely, the genes in the categories "Intracellular Signal Transduction", "Excitability", "GPCR signalling", "Protein cluster", "Neurotransmitter metabolism", "LGIC signalling", "Ion balance/transport", and "G-protein relay" from Table S4 of [Lips et al., 2012] (432 genes in total).
